# Glycogen Synthase Kinase 3 regulates the genesis of the rare displaced ganglion cell retinal subtype

**DOI:** 10.1101/2021.01.06.425300

**Authors:** Elena Kisseleff, Robin J Vigouroux, Catherine Hottin, Sophie Lourdel, Parth Shah, Alain Chédotal, Muriel Perron, Anand Swaroop, Jerome E Roger

## Abstract

Glycogen Synthase Kinase 3 (GSK) proteins (GSK3α and GSK3β) are key mediators of signaling pathways, with crucial roles in coordinating fundamental biological processes during neural development. Here we show that the complete loss of GSK3 signaling in mouse retinal progenitors leads to microphthalmia with broad morphological defects. Both proliferation of retinal progenitors and neuronal differentiation are impaired and result in enhanced cell death. A single wild-type allele of either *Gsk3α* or *Gsk3β* is able to rescue these phenotypes. In this genetic context, all cell types are present with a functional retina. However, we unexpectedly detect a large number of cells in the inner nuclear layer expressing retinal ganglion cell (RGC)-specific markers (called displaced RGCs, dRGCs) when at least one allele of *Gsk3α* is expressed. Excess dRGCs lead to increased number of axons projecting into the ipsilateral medial terminal nucleus, an area of the brain belonging to the non-image-forming visual circuit and poorly targeted by RGCs in wild-type retina. Transcriptome analysis and optomotor response assay suggest that at least a subset of dRGCs in *Gsk3* mutant mice are direction-selective RGCs. Our study thus uncovers a unique role of GSK3 in controlling the genesis of dRGCs, a rare and poorly characterized retinal cell type.

## INTRODUCTION

Glycogen Synthase Kinase 3 alpha (GSK3α) and beta (GSK3β) are functionally redundant serine/threonine kinases encoded by two different genes, sharing 95% identity in their kinase domain (Doble et al., 2007). GSK3 exists at the crossroads of multiple signaling pathways and acts as a key molecular switch to mediate their output and guide distinct cellular processes (Cole, 2012; Doble and Woodgett, 2003; Espinosa et al., 2003; Jin et al., 2009; Shimizu et al., 2008; Wang and Li, 2006). Among the signaling pathways regulated by GSK3 kinases, Wnt canonical pathway is the most well described, with GSK3β inhibition triggering an increase in β-catenin protein levels and its nuclear translocation to activate target gene expression (Doble and Woodgett, 2003).

GSK3 is a key regulator of neural stem/precursor cell proliferation in developing as well as adult brain (Eom and Jope, 2009; Hur and Zhou, 2010; Kim et al., 2009; Pachenari et al., 2017). Conditional deletion and gain of function experiments indicate that GSK3 promotes neuronal differentiation (Hur and Zhou, 2010; Kim et al., 2009). GSK3 exerts its effects through phosphorylation of keys proteins involved in neural development, including proneural factors such as Neurogenin 2 and NeuroD (Li et al., 2012; Moore et al., 2002). In addition, GSK3 fine-tunes the balance between cell death and survival, and its altered function is associated with neurodegenerative pathologies including Alzheimer’s disease, bipolar disorders, and Parkinson’s disease (Golpich et al., 2015; Jacobs et al., 2012; Kremer, 2011; Li et al., 2014; Maurer et al., 2014; Medina et al., 2011).

GSK3 kinases are widely expressed in the developing retina (Pérezleón et al., 2013). GSK3-dependent phosphorylation is shown to control the timing of proneural factor activity and thereby regulate retinal cell fate determination. For instance, inhibition of GSK3 signaling in the developing *Xenopus* retina leads to increase in early-born cell types at the expense of late-born cells (Marcus et al., 1998; Moore et al., 2002).

To elucidate GSK3 function in mammalian retina development, we generated conditional loss-of-function alleles of *Gsk3α* and *Gsk3β* in retinal progenitor cells. We show that complete loss of both GSK3 kinases severely impacts retinal morphology with microphthalmia phenotype, which could be completely rescued with the expression of just one *Gsk3α* or *Gsk3β* wild-type allele. We also noted the presence of a large number of displaced retinal ganglion cells (dRGCs) in the inner nuclear layer in the absence of either *Gsk3α* or *Gsk3β.* In normal conditions, this is a rare retinal cell subtype, poorly characterized so far. Anterograde labeling of the axonal ganglion cell projections into the brain of *Gsk3* mutant mice, allowed us to further support their dRGCs identity. Our study thus identifies GSK3 as a possible determinant of dRGC genesis. We also provide transcriptomic data and visual tests suggesting that at least a subset of these supernumerary dRGCs in *Gsk3* mutant retinas are direction-selective RGCs.

## RESULTS

### Retinal progenitor-specific deletion of both *Gsk3α* and *Gsk3β* results in microphthalmia

We crossed the floxed *Gsk3α^f/f^β^f/f^* mice with α-Cre (αPax6-Cre) line to generate *Gsk3α^f/f^β^f/f^;α-Cre* mice in which *Gsk*3 deletion occurs only in retinal progenitors as early as E10.5 (Marquardt et al., 2001). We first validated our model by assessing the efficacy of *Gsk3α* and *Gsk3β* deletion at E12.5 (Figure 1A). Immunohistochemistry (IHC) using an antibody recognizing both GSK3 proteins showed ubiquitous expression in control retinas (Figure 1A). Both *Gsk3* genes were efficiently deleted in the peripheral retina of *Gsk3α^f/f^β^f/f^;α-Cre* mice, but their expression in the central retina remained preserved consistent with the previously described *α-Cre* expression pattern (Marquardt et al., 2001).

**Figure 1.**
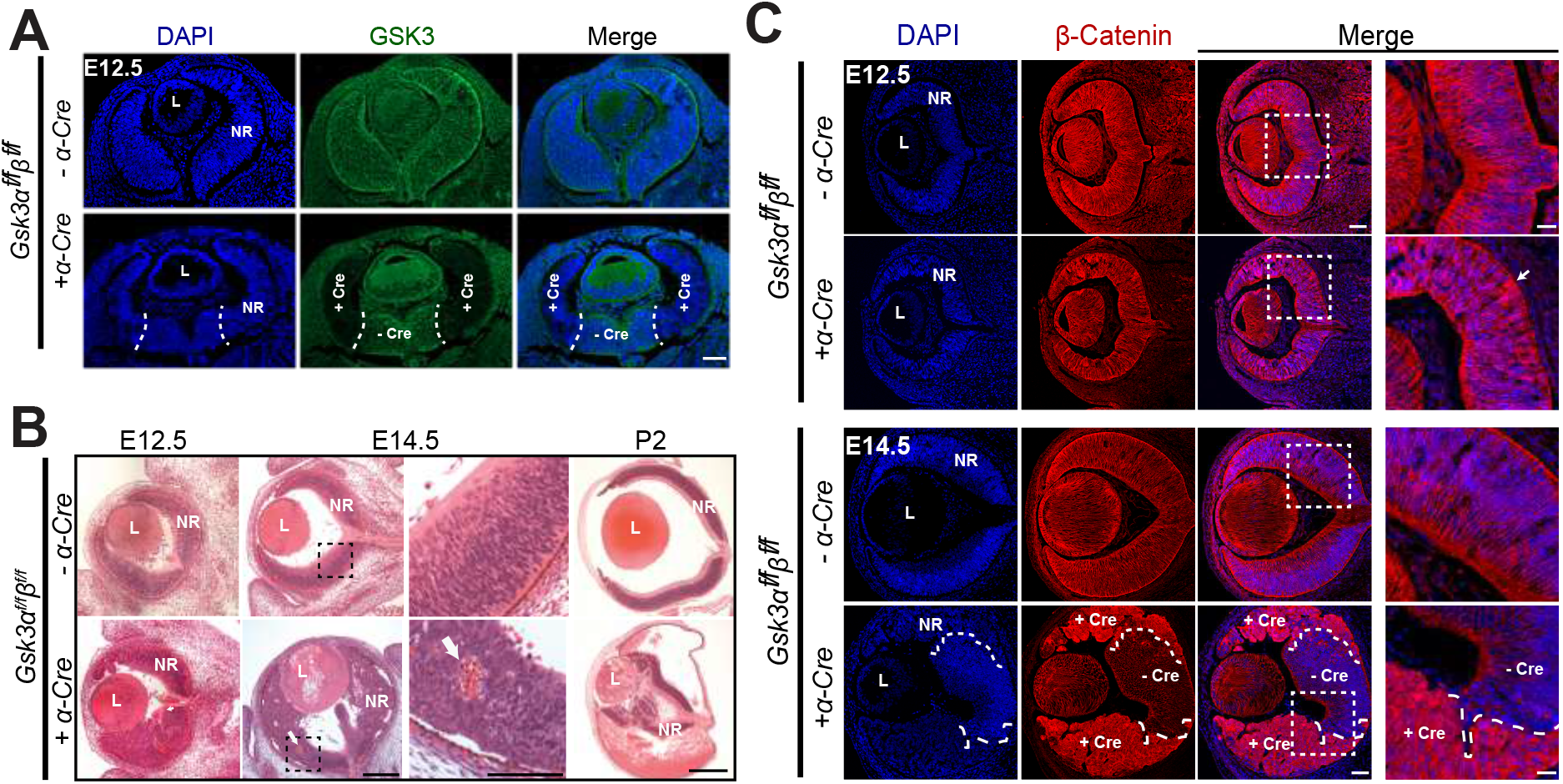
Developmental defects and microphthalmia in *Gsk3*-deficient retina with aberrant nuclear translocation of β-catenin, a key effector of the Wnt canonical pathway. (A) Immunohistochemistry (IHC) of E12.5 retina from *Gsk3α^f/f^β^f/f^* mice expressing or not *α-Cre* using a pan-GSK3 antibody (green) shows efficient deletion at the periphery where the Cre expression has been previously reported (delimited by dashed-line) Scale bar: 100 μm. (B) Hematoxylin and eosin (H&E) staining on methacrylate sections at E12.5, E14.5 and P2 reveals large retinal morphogenesis defects in *Gsk3α^f/f^β^f/f^*;*α-Cre* with blood invasion into the eyeball (showed by white arrow). L, Lens; NR, neural retina. Scale bar: 100 μm at E12.5 and E14.5. 500 μm at P2. For the magnification of P14.5, scale bar: 50 μm. (C) β-catenin accumulation (red) at E14.5 in the Cre-expressing area of *Gsk3α^f/f^β^f/f^*;*α-Cre* animals. White arrow indicates a small area of accumulation. Magnification on the right panel shows the squared delimited area of β-catenin accumulation. L, Lens; NR, neural retina; + Cre, Cre-expressing area; - Cre, area without Cre expression Scale bar: 100 μm, magnification area scale bar: 40 μm.

Hematoxylin and Eosin (H&E) staining revealed major morphological defects with profound retinal disorganization, including the loss of radial arrangement as well as folds and aggregates of retinal progenitor cells (RPCs), in *Gsk3α^f/f^β^f/f^;α-Cre* retina as early as E12.5 (Figure 1B). In addition, blood was detected inside the retinal neuroblastic layer. Structure of the retina worsened rapidly during development although the central part remained unperturbed, consistent with continued *Gsk3* expression in this region. At and after E14.5, the retina was largely reduced whereas the eye size itself was comparable to littermate controls (Figure 1B). A large quantity of blood accumulated inside the eyeball at P2. Finally, growth of the eyeball was severely reduced leading to microphthalmia in the adult (data not shown).

Because of a severe phenotype in the adult, we focused our histological analyses to early retinal development. We first investigated subcellular localization of β-catenin, an established GSK3 target that activates the canonical Wnt pathway (Doble and Woodgett, 2003). In control retina, β-catenin expression was mostly cytoplasmic (Figure 1C). Upon *Gsk3α* and *Gsk3β* deletion, β-catenin was still expressed in the cytoplasm but translocated to the nucleus in some cells as early as E12.5. *Gsk3α^f/f^β^f/f^;α-Cre* retina at E14.5 revealed a clearly defined boundary between the Cre-negative region (GSK3-positive cells) at the center and the Cre-positive region (GSK3-negative cells) at the periphery with nuclear translocation and large accumulation of β-catenin in the nucleus (Figure 1C). Such an expression pattern is consistent with strong activation of canonical Wnt pathway in the absence of both *Gsk3α* and *Gsk3β*. These findings demonstrate that GSK3 regulates Wnt signaling in RPCs and is essential for proper retinal development.

### Lack of both *Gsk3α* and *Gsk3β* in RPCs leads to cell cycle aberrations and retinal progenitor gene deregulation

We focused our analysis on early retinal development (E12.5 and E14.5) in *Gsk3α^f/f^β^f/f^;α-Cre* mice to elucidate the cause of microphthalmia phenotype. To investigate whether proliferation was altered in *Gsk3α^f/f^β^f/f^;α-Cre* mouse retina, a single dose of EdU (to label RPCs in the S-phase) was injected 16 hours prior to harvesting embryos at E12.5 and E14.5. As predicted, EdU-positive cells increased significantly from E12.5 to E14.5 in control retina (Figure 2A, B); yet, no change was detected in *Gsk3α^f/f^β^f/f^;α-Cre* retina and reduction in RPCs in S-phase was evident at E14.5 compared to controls, suggesting proliferation defects. We further examined cell proliferation by labeling late-G2/M-phase retinal progenitors using phospho-histone 3 (pH3) antibody. In control, strongly labeled pH3-positive cells were positioned in the apical surface of the retina where mitosis occurs (M-phase cells). Cells with less intense labeling, presumably late-G2 cells, were present in the outer part of the retinal neuroblastic layer. In *Gsk3α^f/f^β^f/f^;α-Cre* retina, pH3-positive cells located at the periphery were positioned distant from the apical side at both stages (Figure 2A). Moreover, their number at the periphery was abnormal compared to controls, being increased at E12.5 and decreased at E14.5 (Figure 2B). Double pH3- and EdU-positive cells were increased in *Gsk3α^f/f^β^f/f^;α-Cre* retinas compared to controls, at both stages, indicating aberrant cell cycle kinetics (Figure 2B). To investigate the consequence of such cell proliferation defects on the pool of RPCs in the absence of GSK3, we analyzed the expression of Hes1, a retinal progenitor marker (Wall et al., 2009). At E12.5 and E14.5, anti-Hes1 antibody labeled all RPCs throughout control retina with the exception of the basal side where differentiating ganglion cells are located (Figure S1). In *Gsk3α^f/f^β^f/f^;α-Cre* retinas, Hes1 expression was similar to controls at E12.5 indicating the maintenance of RPC pools (Figure S1A). However, by E14.5, Hes1 positive cells were sparser at the periphery where *Gsk3* expression was absent (Figure S1B). In the same area lacking *Gsk3* expression, other progenitor markers, such as Pax6 and Sox2, were similarly decreased in *Gsk3α^f/f^β^f/f^;α-Cre* retina compared to controls (data not shown). Thus, our results show that GSK3 kinases are required for retinal progenitor proliferation and the maintenance of the pool of RPCs.

**Figure 2.**
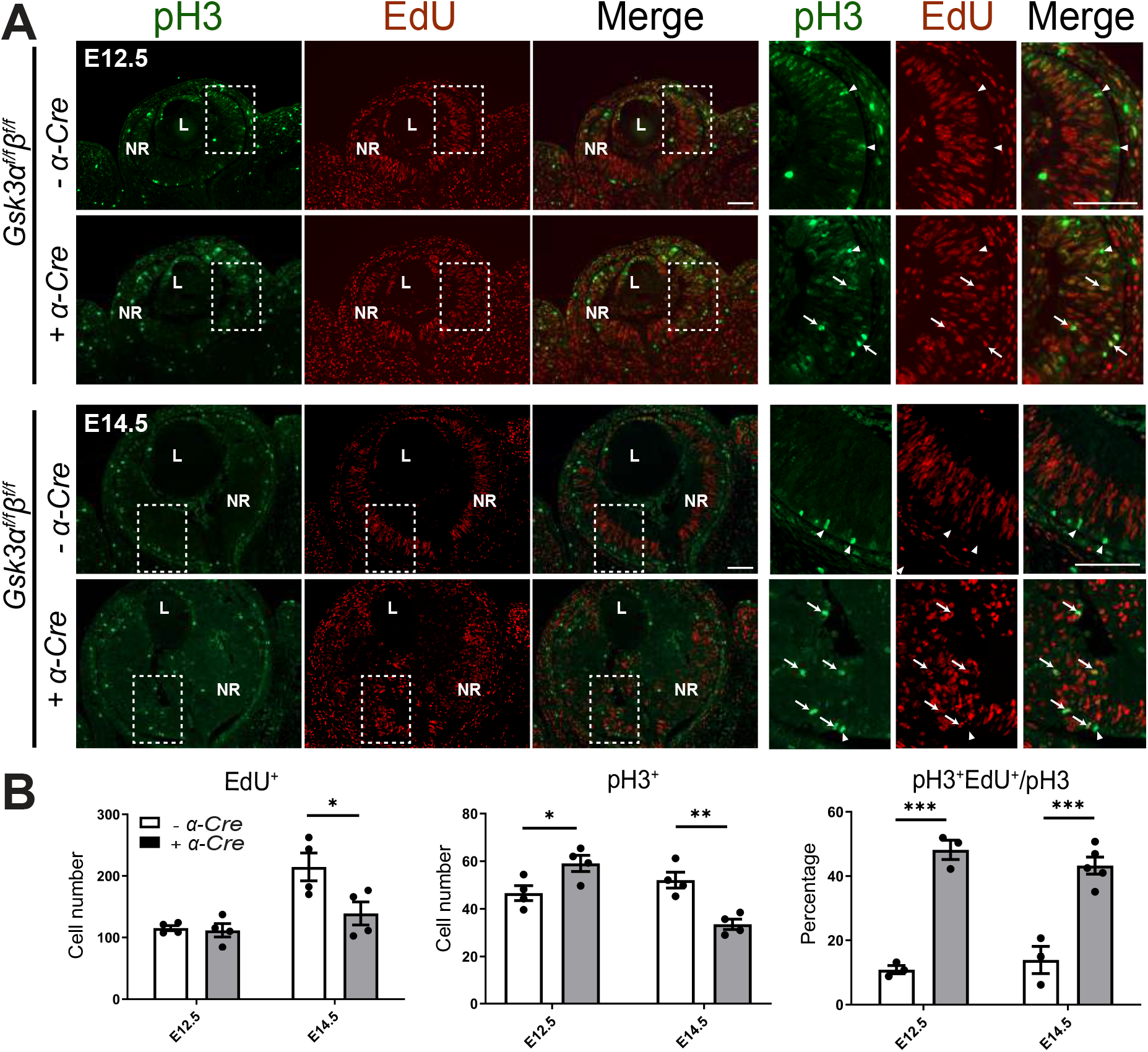
Lack of GSK3 signaling alters cell cycle progression of retinal progenitor cells. (A) Mislocalization of pH3-positive cells in the neuroblastic layer of *Gsk3α^f/f^β^f/f^*;*α-Cre* retina. E12.5 and E14.5 retina stained for pH3 (late-G2/M-phase, green) and EdU (marker of S-phase, red) following a 16H chase. Magnification on the right panel shows the squared delimited area. Arrowheads, pH3-positive and EdU-negative cells; Arrows, pH3- and EdU-positive cells. Scale bar: 100 μm, magnification area scale bar: 100 μm. (B) Lack of GSK3 signaling alters cell cycle progression of retinal progenitors. Quantification at E12.5 and E14.5 of the number of EdU-positive cells (left panel), pH3-positive cells (middle panel) and double positive cells for EdU and pH3 among pH3-positive cells (right panel) in *Gsk3α^f/f^β^f/f^*;*α-Cre* animals and controls. Mean ± SEM values are presented from 3 to 5 independent retinas for each genotype, * indicates *P* ≤ 0.05, ** indicates *P* ≤ 0.01, *** indicates *P* ≤ 0.001.

### Loss of *Gsk3α* and *Gsk3β* impairs retinal progenitor differentiation leading to cell death

To examine whether neuronal differentiation is impacted by the absence of GSK3, we performed IHC using two neuronal markers, Doublecortin (Dcx) and Brn3a, to label newly generated neurons and RGCs, respectively (Figure 3). At E12.5 and E14.5, Dcx- and Brn3a-positive cells were localized in the inner part of the neuroblastic layer of the control retina where retinal differentiation occurs first. In contrast, neuronal differentiation was completely abolished at both stages in *Gsk3α^f/f^β^f/f^;α-Cre* retina, except in the most central part corresponding to the area that does not express Cre recombinase. The absence of retinal differentiation and decreased expression of proliferation markers prompted us to investigate whether the loss of *Gsk3* in RPCs could trigger cell death. We indeed detected a significant increase in the number of TUNEL-positive cells in *Gsk3α^f/f^β^f/f^;α-Cre* retina at both embryonic stages (Figure S2). Thus, microphthalmia observed in the absence of both *Gsk3α and Gsk3β* is likely due to a succession of developmental defects ranging from impaired proliferation and differentiation of RPCs to increased cell death.

**Figure 3.**
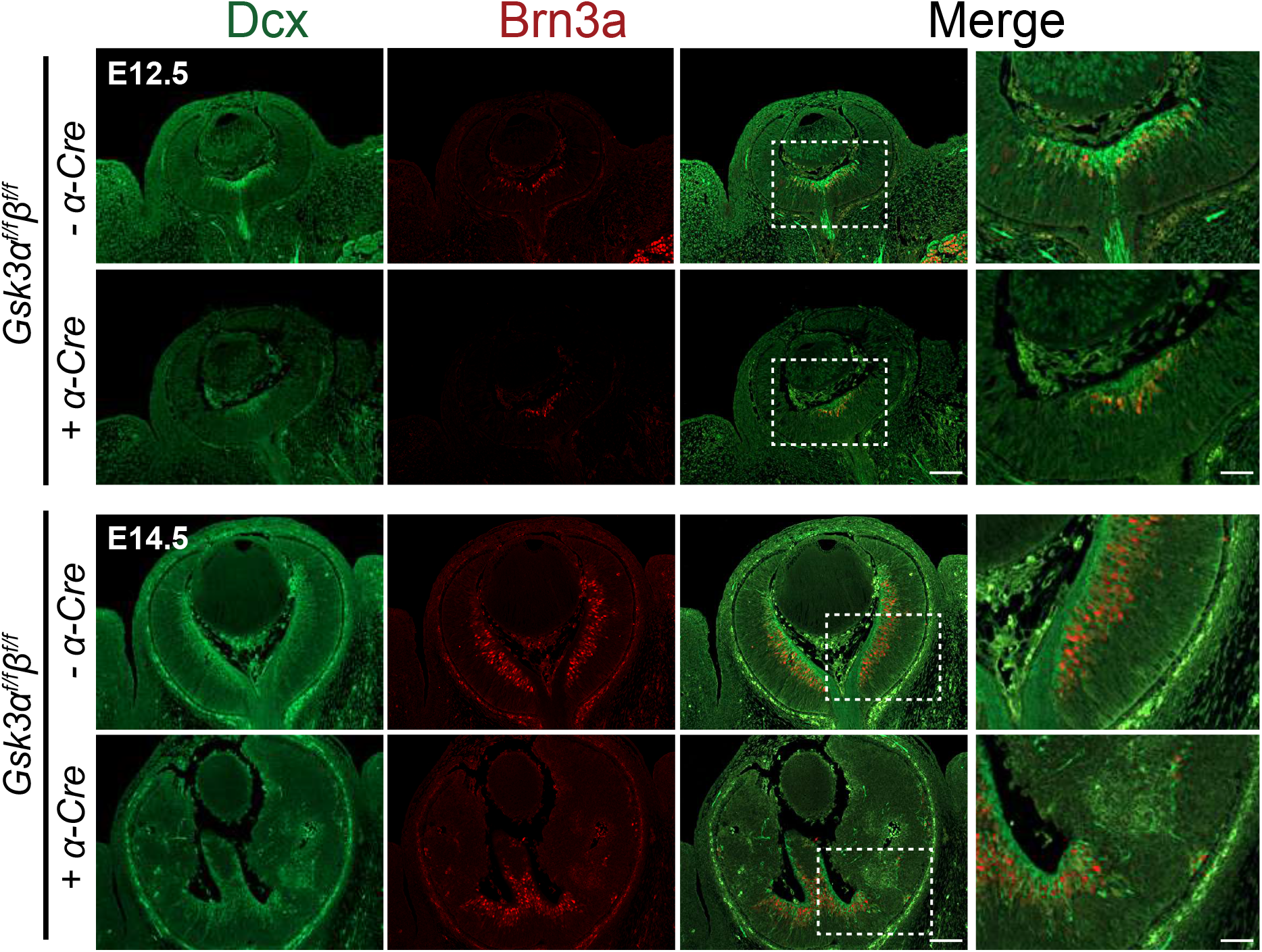
Lack of *Gsk3α* and *Gsk3β* expression in retinal progenitors blocks retinal progenitor differentiation. IHC using Doublecortin (Dcx) (green) and Brn3a (red) antibodies shows the localization of neuronal precursor cells and RGCs, respectively, at the central and basal part of the neuroblastic layer of E12.5 control retina. In contrast, neural differentiation is not observed in *Gsk3α^f/f^β^f/f^*;*α-Cre* retina, with the exception of a small central area where Cre is not expressed. At E14.5, Dcx- and Brn3a-positive cells are distributed across the whole retina in controls, whereas they are absent in periphery in *Gsk3α^f/f^β^f/f^*;*α-Cre* mice. Magnification on the right-hand side shows the squared delimited area and depicts the boundary between a Cre-positive and a Cre-negative area. Scale bar: 100 μm, magnification 40 μm.

### Multiple allelic combinations revealed functional redundancy of *Gsk3α* and *Gsk3β* in retinal development

Severe deleterious effect of the loss of both *Gsk3α and Gsk3β* in RPCs in early development precludes the analysis of late retinal histogenesis. To circumvent this, we generated animals with different combinations of *Gsk3* deletion (loss of only one *Gsk3* gene: *Gsk3α^f/f^β^+/+^;α-Cre* or *Gsk3α^+/+^β^f/f^;α-Cre*, or ¾ deletion: *Gsk3α^f/f^β^f/+^;α-Cre* or *Gsk3α^f/+^β^f/f^;α-Cre*). Immunoblot analysis using anti-GSK3 antibody (recognizing both proteins) in 2-month-old animals with different combinations of *Gsk3α* and *Gsk3β* floxed alleles demonstrated the efficacy of *Gsk3α* and *Gsk3β* deletion (Figure S3A). IHC analysis using anti-GSK3β showed ubiquitous expression of *Gsk3β* in adult control retina and its complete loss in *Gsk3α^f/+^β^f/f^;α-Cre* retina (Figure S3B). At 2-months, retinal histology revealed the correct laminated architecture with normal photoreceptors and interneurons when even one allele of *Gsk3α* (Figure S3C, D) or *Gsk3β* (data not shown) was present. Photopic and scotopic electroretinogram (ERG) recordings, corresponding to cone and rod function respectively, did not show any significant difference between *Gsk3α^f/+^β^f/f^;α-Cre* and control retina (Figure S3E, S3F). These results were similar in mice carrying any combination of *Gsk3* deletion (data not shown). We therefore conclude that a single allele of wild-type *Gsk3α* or *Gsk3β* is sufficient to rescue obvious structural and functional defects in the complete absence of GSK3 signaling.

### Loss of either *Gsk3α* or *Gsk3β* in RPCs leads to increased number of displaced retinal ganglion cells

Even though a single allele of either *Gsk3α* or *Gsk3β* permitted normal retinal development (Figure S3), we observed a striking increase in the number of RGCs, as indicated by Brn3a-positive cells, in the inner nuclear layer (INL) of *Gsk3α^f/+^β^f/f^;α-Cre* retina compared to controls (Figure 4A). Brn3a-positive cells in the INL have been described as displaced retinal ganglion cells (dRGCs), a rare cell type in the mammalian retina (Galli-Resta and Ensini, 1996; Young, 1984). All Brn3a-positive cells in the INL of *Gsk3α^f/+^β^f/f^;α-Cre* retina also expressed NF68 that labels cell bodies and axons of RGCs (Figure 4A). Increased dRGCs were observed in retinas carrying any combination of *Gsk3* deletions tested (*Gsk3α^f/f^β^+/+^, Gsk3α^+/+^β^f/f^*, *Gsk3α^f/+^β^f/f^* or *Gsk3α^f/f^β^f/+^*), with the highest number detected in *Gsk3α^f/+^β^f/f^;α-Cre* mice compared to controls (10-fold increase) (Figure 4B). Interestingly, increase in dRGCs number is not associated with reduced number of RGCs in the GCL, referred to as orthotopic RGCs (oRGCs) (Figure 4B). To validate that these Brn3a-positive cells in the INL were indeed RGCs with axonal projections included in the optic nerve, we performed retrograde labeling with Rhodamine-Dextran applied onto the optic nerve of *Gsk3α^f/+^β^f/f^;α-Cre* mice. Subsequent 3D reconstructions on flat mount retinas revealed the presence of numerous fluorescent cell bodies located in the INL compared to controls, demonstrating that axons of dRGCs indeed reached the optic nerve (Figure 4C). Thus, Brn3a-positive cells located in the INL of *Gsk3α^f/+^β^f/f^;α-Cre* retinas are indeed RGCs.

**Figure 4.**
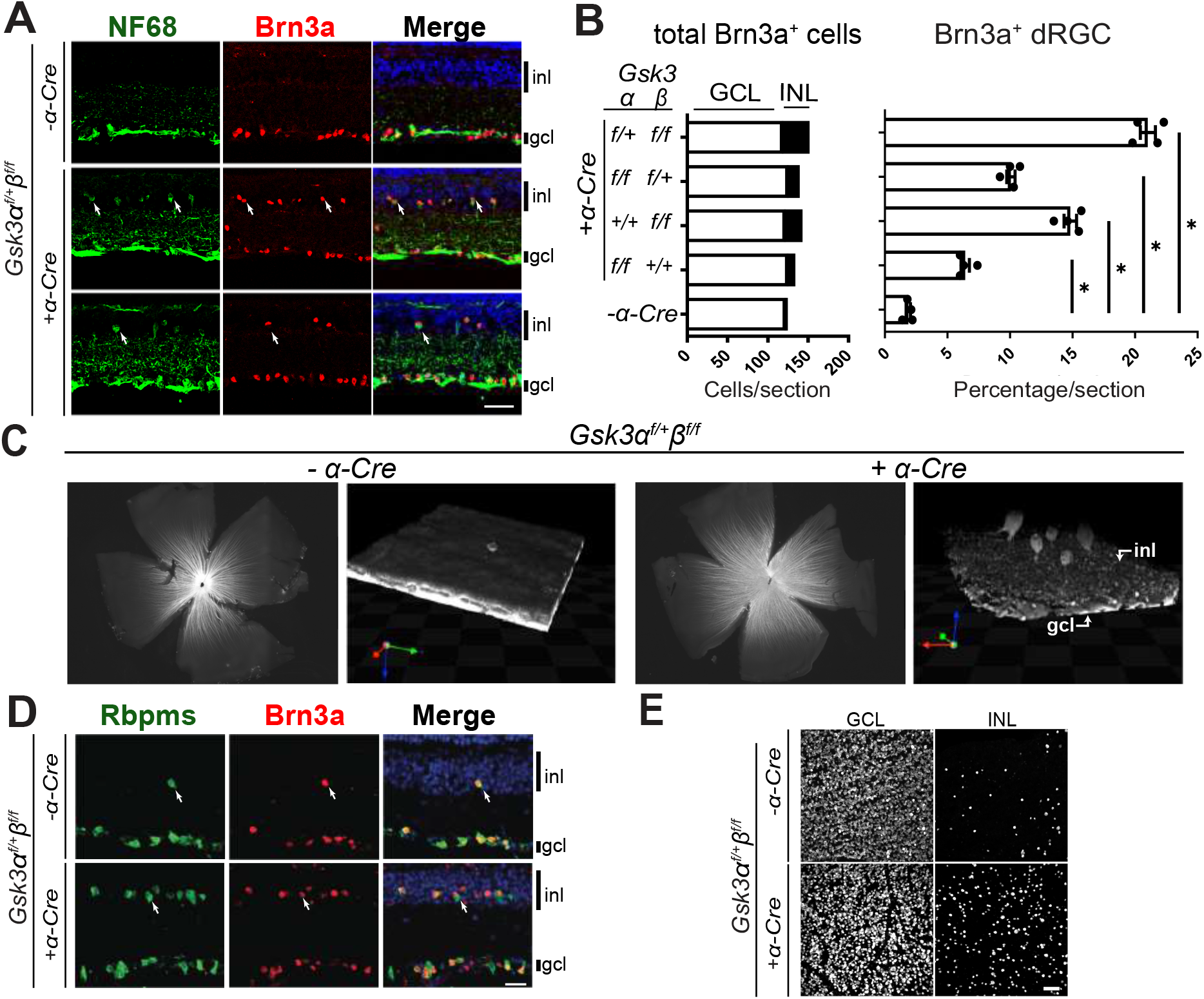
Gradual loss of *Gsk3α* and/or *Gsk3β* leads to an increased number of Brn3a-positive retinal ganglion cells displaced in the inner nuclear layer (INL) of adult retina. (A) Brn3a (red) and NF68 (green) IHC on 2-month-old *Gsk3α^f/+^β^f/f^; α-Cre* mouse retina reveals the presence of supernumerary displaced retinal ganglion cells (dRGCs, arrows) in the INL of *Gsk3α^f/+^β^f/f^; α-Cre* compared to littermate controls. Top panel represents control retinas, middle panel a peripheral retinal area, and bottom panel a more central area. Scale bar: 20 μm. (B) Gradual loss of *Gsk3α* and *Gsk3β* alleles (*Gsk3α^f/f^β^+/+^, Gsk3α^+/+^β^f/f^*, *Gsk3α^f/+^β^f/f^* or *Gsk3α^f/f^β^f/+^*) leads to a gradual increase of Brn3a-positive RGCs located to the INL, with the highest number observed in *Gsk3α^f/+^β^f/f^*; *α-Cre* animals. Left stacked histogram represents counting of the total number of Brn3a-positive cells per section located in the GCL (white bars) and in the INL (black bars). Right histogram represents the percentage of the dRGCs among the total number of Brn3a-positive cells per section for each combination. Mean ± SEM values are presented from 4 biological replicates, * indicates *P* ≤ 0.05. (C) dRGCs send their axons into the optic nerve. Visualization of dRGCs after 3D reconstruction of 2-month-old flat mounted retina of control and *Gsk3α^f/+^β^f/f^*; *α-Cre* animals following retrograde labeling with Rhodamin-Dextran applied onto the optic nerve. inl: inner nuclear layer, gcl: ganglion cell layer. (D) Brn3a (red) and Rbpms (green) IHC on 2-month-old mouse retina reveal the co-expression of these two RGC markers (dRGCs, arrows) in the INL of both *Gsk3α^f/+^β^f/f^; α-Cre* dRGCs and in littermate controls. Scale bar: 20 μm. (E) Flat mounted retina from *Gsk3α^f/+^β^f/f^; α-Cre* and littermate controls labelled with anti-Rbpms antibody demonstrated the large number of Rbpms-positive dRGCs in the INL of *Gsk3α^f/+^β^f/f^; α-Cre* mice.

Due to their low number in control retina (around 2%), dRGCs have been poorly characterized with very few markers identified, such as Brn3a (Nadal-Nicolás et al., 2012; Nadal-NicolÃ¡s et al., 2014). Immunostaining on sections and flat mount retinas with additional RGC marker antibodies revealed that dRGCs in *Gsk3α^f/+^β^f/f^;α-Cre* retina were also positive for Rbpms (Rodriguez et al., 2014) confirming their increased number in the INL compared to controls (Figure 4D, E). Similar results were observed with Islet1 labeling (Figure S4) (Bejarano-Escobar et al., 2015). Finally, Brn3a-positive dRGCs did not express markers of other INL neurons such as Choline-Acetyltransferase (CHAT, amacrine cells), Calretinin (amacrine cells), or Calbindin (horizontal cells) further confirming their RGC identity (data not shown).

To test whether dRGCs in *Gsk3* mutant mice were produced during the same developmental time window as oRGCs, we performed pulse chase experiments by injecting EdU at E12.5, at the peak of RGC birth. Retinal sections from one-month-old animals were then immunolabelled using anti-Brn3a antibody (Figure 5A). In control and *Gsk3α^f/+^β^f/f^*;*α-Cre* retina, we identified 40-50% of RGCs that were Brn3a/EdU-positive in all layers examined (GCL and INL), indicating that both dRGCs and oRGCs were born around the same time (Figure 5B). We next examined whether dRGCs are usually overproduced during normal retinal development and eliminated later on. In this context, increased number of dRGCs in *Gsk3α^f/+^β^f/f^*;*α-Cre* retinas could result from a defect in dRGC riddance occurring during the first two postnatal weeks, a period of developmental cell death in the retina (Young, 1984). At P0, the number of Brn3a-positive oRGCs was similar between littermate control and *Gsk3α^f/+^β^f/f^*;*α-Cre* retina (Figure 5C and D). In contrast, the proportion of Brn3a-positive cells located in the inner part of neuroblastic layer corresponding presumably to dRGCs was much lower in control retinas (6±0.1%) compared to *Gsk3α^f/+^β^f/f^*;*α-Cre* retinas (30±1.4%). Thus, dRGCs are not overproduced and eliminated postnatally in control retina. Our results demonstrate that dRGCs are generated during early waves of retinogenesis in *Gsk3α^f/+^β^f/f^*;*α-Cre* retina and strongly suggest that GSK3 kinases play a role in restricting their number during normal retinal development.

**Figure 5.**
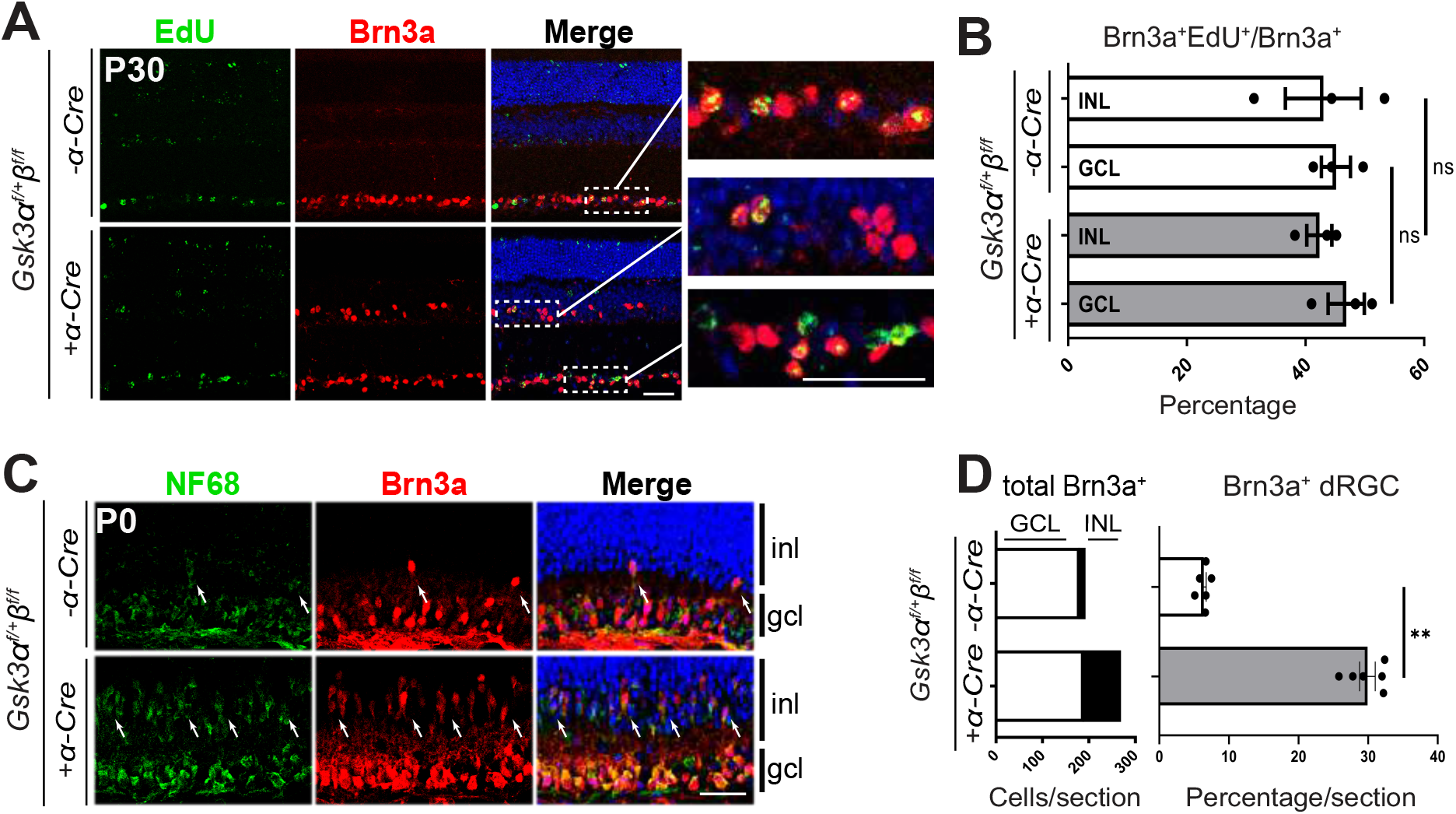
dRGCs are produced in the same differentiation wave as oRGC located in the GCL. (A) EdU- (green) and Brn3a-positive cells (red) were found both in the GCL and in the INL of 30-days old *Gsk3α^f/+^β^f/f^*; *α-Cre* animals after a single injection of EdU at E12.5. (B) Percentage of EdU- and Brn3a-positive cells located either in the GCL or in the INL among total number of Brn3a-positive cells. Mean ± SEM values are presented from 3-4 biological replicates, ns: not significant (C) Brn3a (*red*) and NF68 (green) immunostaining on P0 mouse retina revealed that a large number of dRGCs were already present in *Gsk3α^f/+^β^f/f^; α-Cre* but they were fewer in littermate controls (white arrows). (D) Left stacked histogram represents counting of the total number of Brn3a-positive cells per section located in the GCL (white bars) and in the INL (black bars) of *Gsk3α^f/+^β^f/f^; α-Cre* retina. Right histogram represents the percentage of the dRGCs among the total number of Brn3a-positive cells per section. Mean ± SEM values are presented from 6 biological replicates, ** indicates *P* ≤ 0.01. inl, inner nuclear layer; gcl, ganglion cell layer. Scale bar: 20μm.

### dRGCs produced in the absence of either *Gsk3α* or *Gsk3β* project to accessory visual system circuitry

Previous studies in birds and reptiles have reported that dRGCs could be responsible for optokinetic nystagmus, as they mostly project to the accessory optic nuclei (AOS) (Cook and Podugolnikova, 2001), which is critical for non-image forming circuit and image stabilization (Simpson, 1984). To test whether dRGCs in *Gsk3* mutants project into specific visual nuclei in the brain, including the AOS, we traced the total pool of RGCs, including dRGCs, with Cholera Toxin beta subunit (CTB). Bilateral injection of CTB, coupled to either an alexa-555 or -647 followed by 3D imaging, allowed us to trace both ipsi- and contra-lateral projecting axons. We first confirmed that CTB injections indeed marked the dRGCs based on flat mounts of retinas after Brn3a immunolabelling (data not shown). To visualize the entire visual projection network, we carried out whole-brain clearing using iDISCO+ followed by light-sheet fluorescent imaging and 3D reconstruction (Figure 6A). Complete loss of *Gsk3β* displayed a specific increase in ipsilateral projecting RGCs specifically in one of the three nuclei composing the AOS, the Medial Terminal Nucleus (MTN) (Simpson, 1984). This terminal nucleus is the main component of the AOS reacting best to either upward or downward movement and mediates the optokinetic nystagmus critical for image stabilization (Yonehara et al., 2009). Calculation of the signal intensity ratio between the ipsilateral and contralateral MTN demonstrated a significant increase of RGC projections into the ipsilateral MTN in retinas with *Gsk3β* deletion (Figure 6B). This result strongly suggests that excess dRGCs might participate to the non-image forming circuit.

**Figure 6.**
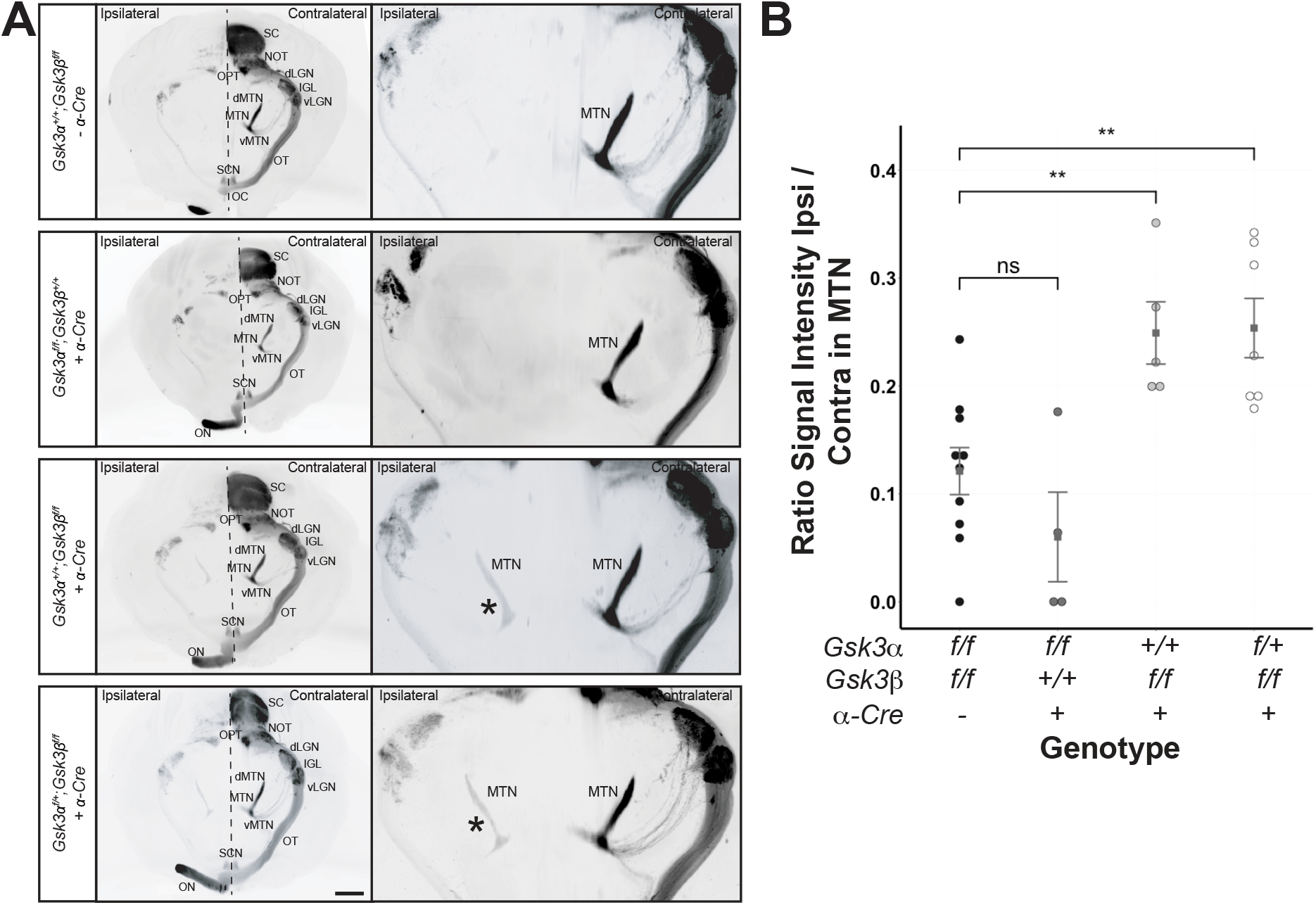
Lack of Gsk3β results in RGC projections into the ipsilateral Medial Terminal Nucleus. (A) All panels are light sheet fluorescence microscopy of solvent-cleared adult brain from control, *Gsk3α^f/f^β^+/+^; α-Cre*, *Gsk3α^+/+^β^f/f^; α-Cre* and *Gsk3α^f/+^β^f/f^; α-Cre* animals after intravitreal injection of CTB coupled to either an Alexa-555 or -647. Ipsilateral projections of RGCs into the MTN was observed in the absence of *Gsk3β* expression. SC, superior colliculus; NOT, nucleus of optic tract; dLGN, dorsal lateral geniculate nucleus; vLGN, ventral lateral geniculate nucleus; IGL, intergeniculate leaflet; OPT, Olivary Pretectal Nucleus; dMTN, dorsal medial terminal nucleus; MTN, medial terminal nucleus; vMTN, ventral medial terminal nucleus; OT, Optic tract; SCN, suprachiasmatic nucleus; ON, optic nerve. Scale bar: 1mm; * indicates the ipsilateral MTN. (B) Quantification of the signal intensity ratio between ipsilateral and contralateral MTN in controls and *Gsk3* mutants (including *Gsk3α^f/f^β^+/+^; α-Cre*, *Gsk3α^+/+^β^f/f^; α-Cre*, and *Gsk3α^f/+^β^f/f^; α-Cre*). ns: non-significant, ** *P* ≤ 0.01.

### Whole transcriptome analysis suggests that dRGCs in GSK3 mutant retinas are direction-selective ganglion cells

We next performed transcriptome analysis using RNA-sequencing in order to identify molecular changes in adult *Gsk3α^f/+^β^f/f^*;*α-Cre* retina and to better characterize dRGCs. Retinas from *Gsk3α^f/+^β^f/f^* mice were used as controls. Gene level analysis revealed 111 differentially expressed genes (DEGs) using filtering criteria of Fold Change (FC) = 1.5 with a False Discovery rate (FDR) cutoff of ≤ 0.05 and a minimum mean expression value of one FPKM (fragments per kilobase of exon per million reads mapped) in at least one of the two experimental groups (Figure 7A and Figure S5). Pathway analysis of DEGs revealed several statistically significant overrepresented pathways (Figure S6). Biological processes and molecular functions pathways included 48 DEGs; of these, 33 genes were expressed in RGCs based on published whole transcriptomic data from purified RGCs (for a total 69 RGC-expressed genes among the 111 DEGs) (labeled with stars in Figure 7B) (Sajgo et al., 2017). Dominance of RGC-expressed genes in our dataset is consistent with the high number of dRGCs observed in *Gsk3α^f/+^β^f/f^*;*α-Cre* retina.

**Figure 7.**
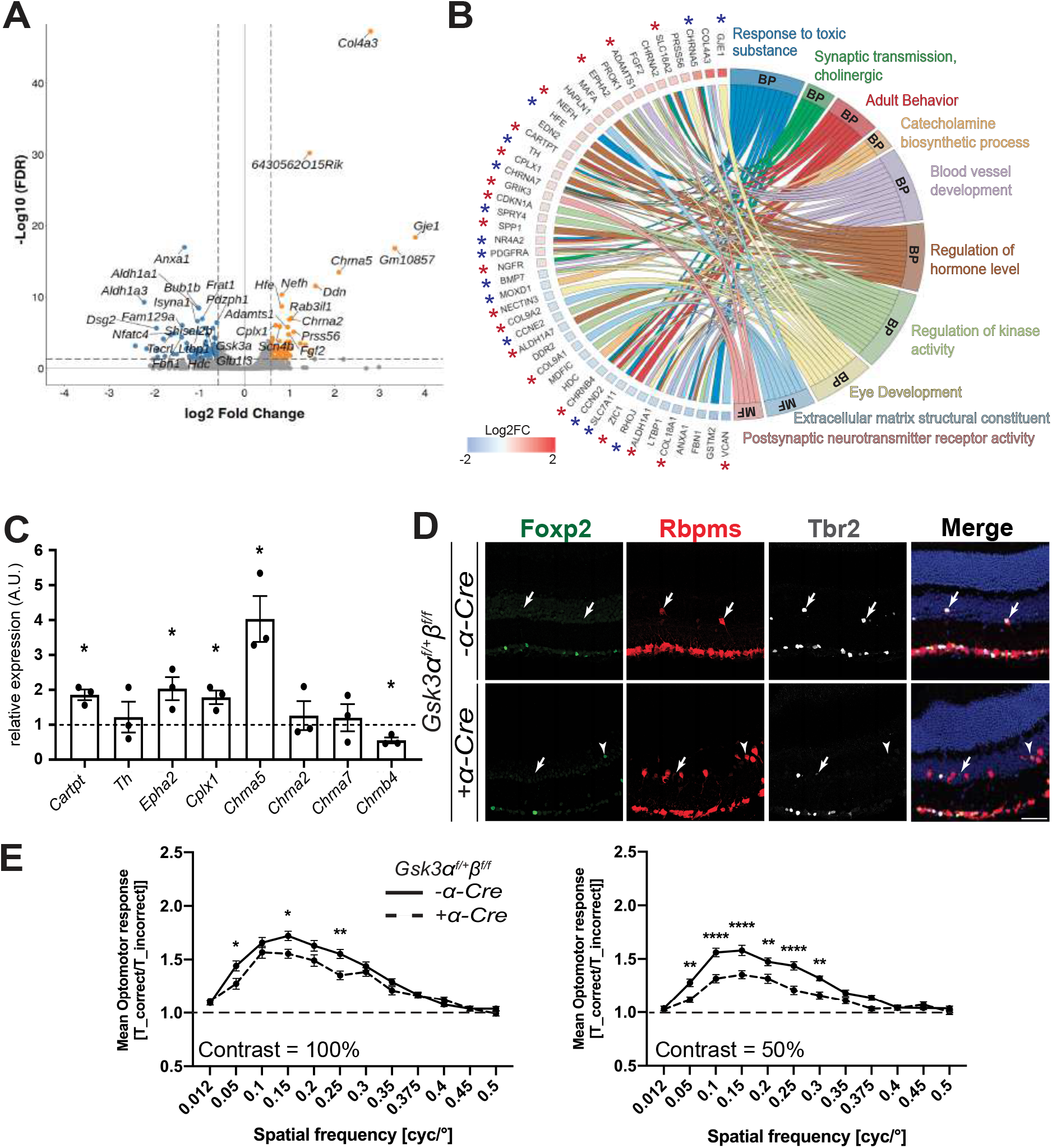
dRGC characterization in *Gsk3α^f/+^β^f/f^*; *α-Cre* retina. (A) Volcano plot representation of differentially expressed genes between *Gsk3αf/+βf/f; α-Cre* and control retina plotted on the x-axis (log2 scale). FDR adjusted significance is plotted on the y-axis. Orange and blue dots: significantly up-regulated and down-regulated genes in *Gsk3α^f/+^β^f/f^; α-Cre* retina, respectively. Vertical dashed lines represent FC=1.5. Horizontal dashed line represents FDR=0.05. (B) Chord plot representation of DEGs related to GO annotations belonging to either molecular functions (MF) or biological process (BP). Overlaps in GO annotation amongst genes within each category are visualized. *\** correspond to genes expressed in previously published purified RGCs (blue, slightly expressed genes in RGCs between 1 and 5 FPKM; red, highly expressed genes in RGCs more than 5 FPKM). (C) RT-qPCR validation of selected DEGs identified by RNA-seq analysis. Differential expression analysis by RT-qPCR of *Cartpt, Th, Epha2, Cplx1, Chrna5, Chrna2, Chrna7, Chrnb4* in *Gsk3α^f/+^β^f/f^; α-Cre* retina at 2-months of age, relative to littermate control retina levels. All values are expressed as the Mean ± SEM from three biological replicates. * indicates *P* ≤ 0.05. (D) IHC on 2-month-old mouse retina reveals the presence of a subset of dRGCs (Rbpms-positive dRGCs, red) in *Gsk3α^f/+^β^f/f^; α-Cre* expressing either the transcription factor Tbr2 (grey) or Foxp2 (green). Arrows indicate Tbr2 and Rbpms-positive dRGCs; arrowheads represent Foxp2 and Rbpms-positive dRGCs. Scale bar: 50 μm. (E) The mean OMR indices (±SEM) are plotted as a function of spatial frequency for each genotype (n=13 for *Gsk3α^f/+^β^f/f^* and 18 for *Gsk3α^f/+^β^f/f^*;*α-Cre* genotype). The baseline (1; dashed line) represents unspecific head movements and no response to the stimulus. OMR at 100% and 50% contrast in *Gsk3α^f/+^β^f/f^; α-Cre* mice (dashed line) and controls (black line). * indicates *P* ≤ 0.05, ** indicates *P* ≤ 0.01, *** indicates *P* ≤ 0.001.

Among interesting candidates dysregulated in the biological processes and molecular function pathways (Figure 7B and S6), we identified *Chrna2*, *Chrna5, Chrna7,* and *Chrnb4*. With the exception of *Chrna2*, all of these are reported in purified ganglion cell transcriptomic data (Sajgo et al., 2017). RT-qPCR analysis validated upregulation of *Chrna5*, as well as the downregulation of *Chrnab4* (Figure 7C). For *Chrna7*, we could only find a trend toward the increase although not significant. We hypothesize that upregulation of *Chrna5,* and potentially that of *Chrna7,* results from the increase in dRGCs in *Gsk3α^f/+^β^f/f^*;*α-Cre* retina. Another gene *Cartpt,* encoding preprotein CART (Cocaine- And Amphetamine-Regulated Transcript Protein) that was upregulated in *Gsk3α^f/+^β^f/f^*;*α-Cre* retina was validated by RT-qPCR (Figure 7C). *Cartpt* is specifically expressed in direction-selective RGCs (DS-RGCs) (Kay et al., 2011; Rousso et al., 2016), suggesting that dRGCs (or at least a subset) in *Gsk3α^f/+^β^f/f^*;*α-Cre* retina might be DS-RGCs. In support with this hypothesis, we found some dRGCs in *Gsk3α^f/+^β^f/f^*;*α-Cre* and littermate control retina positive for the transcription factor Tbr2, described as essential for RGC specification participating in non-image-forming visual circuits (Figure 7D) (Sweeney et al., 2014, 2019). A small subset of dRGCs also expressed Foxp2, a transcription factor involved in DS-RGC differentiation in mice (Figure 7D) (Rousso et al., 2016; Sato et al., 2017). These two factors were expressed in a mutually exclusive way in Rpbms-positive dRGCs, suggesting that dRGCs in *Gsk3α^f/+^β^f/f^*;*α-Cre* might encompass several subtypes.

### Optomotor response is impaired in GSK3 mutant

Given that DS-RGCs are reported to drive the optomotor response (OMR) by projecting mainly into the contralateral AOS (Simpson, 1984; Yonehara et al., 2009), we tested the OMR of *Gsk3α^f/+^β^f/f^*;*α-Cre* mice. The OMR indices (T_correct / T_incorrect) were calculated from three trials at contrast 1 and 0.5 (Figure 7E). At 100% contrast, the OMR indices were significantly reduced in *Gsk3α^f/+^β^f/f^*;*α-Cre* mice compared to controls at 0.05, 0.15 and 0.25 cycles per degree (cpd). The maximum OMR index was observed at 0.15 cpd in controls whereas it reached its maximum at 0.1 in *Gsk3α^f/+^β^f/f^*;*α-Cre* mice. At 50% contrast, the OMR indices were also significantly reduced in *Gsk3α^f/+^β^f/f^*;*α-Cre* mice compared to controls but to a larger extend between 0.05 and 0.3 cpd. The maximum OMR index was observed at 0.15 cpd in both controls and *Gsk3α^f/+^β^f/f^*;*α-Cre* mice. Altogether, these results demonstrate an impaired OMR in *Gsk3α^f/+^β^f/f^*;*α-Cre* mice. These data, in support with our transcriptomic and axonal projections analyses, suggest that at least a subset of dRGCs expressing only one allele of *Gsk3α* are DS-RGCs.

## DISCUSSION

In this study, we report that complete loss of GSK3 in retinal progenitors leads to microphthalmia in adult mice resulting from a successive chain of catastrophic molecular events causing severe morphological defects with progressive death of the pool of proliferative retinal progenitors and lack of neuronal differentiation. Such a severe phenotype was not observed anymore when only one *Gsk3α* or *Gsk3β* allele was expressed, confirming the functional redundancy of these two genes. Our results suggest that the kinase GSK3 is the first discovered dRGCs determinant during retinal histogenesis. We indeed found that mouse retinas with only one allele of *Gsk3* exhibit an excessive number of dRGCs. The concomitant large increase of axonal projections to the ipsilateral MTN, our RNA-Seq data and optomotor response tests, led us to propose that these dRGCs are involved in the detection of image motion direction.

We reported that complete loss of GSK3 activity in retinal progenitors leads a successive chain of catastrophic molecular events causing severe morphological defects with progressive death of the pool of proliferative retinal progenitors and lack of neuronal differentiation. A lack of cell differentiation was also described in the developing brain of *Gsk3* mutant mice (Kim et al., 2009). Surprisingly however, we demonstrate that the absence of GSK3 impairs the maintenance of the pool of retinal progenitors and their survival, leading subsequently to microphthalmia in adults. Our results are in striking contrast with the “big head phenotype” due to a large expansion of the pool of neural progenitors by hyperproliferation and the absence of cell death at early stages observed in *Gsk3α*^−/−^;*Gsk3β^f/f^*; *Nestin-Cre* embryos (Kim et al., 2009). Retinal progenitors might be more prompt to cell death if they encounter proliferative defects compared to brain neural progenitors. GSK3 kinases are established negative regulators of the Wnt canonical pathway (Doble and Woodgett, 2003). As such, we found that lack of GSK3 leads to increased amount of β-catenin nuclear translocation. Interestingly, the effect of *Gsk3* loss of function in the retina mimics the effect of β-catenin gain of function. Indeed, constitutive retinal activation of β-catenin does not elicit an hyperproliferation of retinal cells, as one could have expected from Wnt pathway activation but instead, it decreases cell proliferation and neural differentiation, resulting in a small eye phenotype (Fu et al., 2006; Ouchi et al., 2011). Besides, β-catenin has been recognized as a key regulator of cell adhesion (Ouchi et al., 2011). The lamination defects observed in *Gsk3* knockout mice may thus also result from altered cell adhesion due to β-catenin activation.

In pigmented wild-type mouse retina, dRGCs in the INL are a very rare and poorly-described type of cells representing only 2% of RGCs (Balkema and DrÄger, 1990; Doi et al., 1994; Dräger and Olsen, 1980; Nadal-NicolÃ¡s et al., 2014). It is therefore striking that dRGC number increases up to 20% when a single copy of *Gsk3α* is present in retinal progenitors. To our knowledge, such a high number of dRGCs has never been reported in a transgenic/mutant animal. A previous study hypothesized that dRGCs are misplaced in the INL due to an ontogenic aberration rather than representing an independent class of RGCs (Buhl and Dann, 1988; Doi et al., 1994). Indeed, differential cell adhesion plays a key role in sorting and migration of retinal cells in their appropriate layers, especially for RGCs. One can therefore hypothesize that enhanced dRGCs in mice with a single copy of *Gsk3* is the consequence of increased aberration events. This hypothesis could be supported by our RNA-Seq data showing the upregulation of genes coding for collagen subunits (*Col18a1*, *Col4a3*, *Col9a1*, *Col9a2*) and extracellular matrix proteins in *Gsk3α^f/+^β^f/f^*;*α-Cre* retina, which could favor migration defects. Noticeably, if it were the case, the increase in dRGCs should be accompanied by a decrease in oRGCs. However, we found that the number of oRGC in the GCL is unaltered, strongly suggesting that RGCs in the INL of mice with a single copy of *Gsk3* represent a specific subtype of RGCs. In support with this, topographic and quantitative analysis of RGCs in albinos and pigmented rats indicate that dRGCs are not misplaced by ontogenic mistakes but indeed represent a specific subpopulation of RGCs (Nadal-NicolÃ¡s et al., 2014). GSK3β is involved in neural cell fate decision by controlling the timing of the activity of bHLH transcription factors, such as NeuroD or Neurog2 (Li et al., 2012; Moore et al., 2002). If dRGCs are not produced following ontogenic aberrations but are instead determined by a proper genetic program, it would be interesting to identify transcription factors involved and seek for any regulation by GSK3 kinases.

In reptiles, amphibians and birds, only dRGCs project into the MTN, whereas in mammals only oRGCs have been reported as projecting into the MTN (Fite et al., 1981; Krause et al., 2014). Our results obtained from anterograde labeling clearly demonstrated a large increase in ipsilateral MTN projections in absence of *Gsk3β*, whereas it was absent or very dim in control animals. Although this strongly suggests that excess dRGCs in mutant mice are causing this phenotype, we cannot exclude the possibility that mutant oRGCs also participate to these ipsilateral MTN projections. However, contralateral projections did not seem to be affected. We can speculate that the low number of ipsilateral MTN projections in the control condition reflects the low number of dRGCs present in the WT retina and could therefore explain that such result had not been described so far. Altogether, our results strongly suggest that dRGCs may primarily project into the ipsilateral MTN. In mice, it has been shown by retrograde labeling from the superior colliculus (SC), which receive large amount of RGC projections, that dRGCs/oRGCs project to one or both SCs (Karten et al., 1977). Although challenging, similar experiments, *i.e.* fluorescent dye injection into the ipsilateral MTN, may allow to discriminate whether the increased signal in absence of *Gsk3β* originates only from dRGCs and whether these cells also project into this area in WT retina. In regards to our results, it is still unclear whether GSK3 function is to limit the number of dRGCs and actively regulates their correct projection to the contralateral MTN or if GSK3 function is limited to the tight control of dRGC number, which project thereafter to the ipsilateral MTN on a GSK3-independent manner.

Given the very low percentage of dRGCs in the control retina, their function is poorly studied in mammals. In contrast, dRGC function, projections and topography have been extensively investigated in bird and reptile retina (Mouritsen et al., 2004). In birds, cryptochrome-expressing dRGCs are used as a magnetic compass for orientation (Nießner et al., 2016). In European Robin birds, *Erithacus rubecula,* a low number of dRGCs have been identified but specifically express Cryptochrome 1b only during nocturnal migration period (Nießner et al., 2016). In rodents, retrograde labeling from the optic nerve led to the identification of 16 classes of dRGCs based on their ramification levels of their dendrites as well as the dendritic field size (Pang and Wu, 2011). Based on dRGC dendrite projections into the IPL, it has been proposed that most dRGCs in WT retina are functionally more involved in retinal OFF light pathways (Pang and Wu, 2011). Similar methods applied to *Gsk3α^f/+^β^f/f^*;*α-Cre* retina should shed more light on dRGC function and establish whether all the different classes are present. As part of the AOS, the MTN receives afferent signal from the eye and sends efferent signal to motor neurons controlling the position of the eye. As such, optokinetic reflex relies on direction specific retinal projections to the AOS. Neurons of the dorsal terminal nucleus (DTN) codes for horizontal stimulus whereas neurons of the MTN codes for vertical stimulus (Giolli et al., 2006; Yonehara et al., 2009). Therefore, the direction of image motion relies on DS-RGCs in the retina. The alteration the OMR in *Gsk3α^f/+^β^f/f^*;*α-Cre* mice, support the hypothesis that some of the supernumerary dRGCs are indeed related to motion detection. Although the number of dRGCs was drastically increased, the OMR was not increased but at the contrary reduced. Such result might be caused by the higher number of projections to the ipsilateral side instead of the contralateral one, leading to an alteration of the neuronal circuit regulating the OMR (Simpson, 1984; Yonehara et al., 2009). We also identified in *Gsk3α^f/+^β^f/f^*;*α-Cre* and control retinas a small subset of dRGCs positive for the transcription factors Tbr2 and Foxp2, markers for non-image-forming RGCs and DS-RGCs respectively (Nadal-NicolÃ¡s et al., 2014). Together with our transcriptomic data (upregulation of genes such as *Cartpt* (expressed in DS-RGCs)), these results strongly suggest that the large number of dRGCs in *Gsk3α^f/+^β^f/f^*;*α-Cre* retina might indeed be DS-RGCs projecting into the MTN. It has been recently proposed that dRGCs might be also involved in predator detection by integrating overhead visual information (Nadal-NicolÃ¡s et al., 2014). Using suitable and complementary visual tests, our genetic model with an excess of dRGCs could be highly valuable to complete the functional identification of the dRGCs in vision.

Overall, our results demonstrate the critical role of GSK3 in tightly regulating the number of a rare type of dRGCs, poorly described yet. Therefore, *Gsk3* mutant mice, encompassing a large number of dRGCs in their retina, offer a unique and powerful model system to further study the embryonic origin, synaptic connections and visual function of dRGCs in mammals.

## MATERIALS AND METHODS

### Animals and tissue collection

All animal experiments have been carried out in accordance with the European Communities Council Directive of 22 September 2010 (2010/63/EEC), European Union guidelines effective and with the Association for Research in Vision and Ophthalmology statement for the use of animals in ophthalmic and visual Research. All animal care and experimentation were also conducted in accordance with guidelines, under the license APAFIS#1018-2016072611404304 by the Institutional animal care committee n°059 in France and by Animal Care and Use Committee at the National Institutes of Health (ASP#650). *Gsk3α* and *Gsk3β* floxed mice were generously provided by Dr. Jim Woodgett (University of Toronto, Canada). Floxed *Gsk3* mice were mated with those carrying the retina-specific regulatory element of murine *Pax6* driving the expression of the Cre recombinase (*α-Cre*) in retinal progenitors as early as E10.5 generously provided by Peter Gruss (Marquardt et al., 2001). Mice from either sex were used for experimental procedures. All mouse genotyping was performed as described (Roger et al., 2010).

### Hematoxylin & eosin (H&E) staining, and immunostaining

Methacrylate sections were used for H&E staining as previously described (Roger et al., 2012). For IHC on frozen sections, enucleated eyeballs were fixed at the required stage in 4% PFA for 60 min on ice and incubated in an increasing concentration of sucrose (10%, 20% and 30%), then embedded in OCT. Embedded eyeballs were serially cut to 12 μm sections using a cryostat. For embryonic stages, pregnant females were euthanized and whole heads of pups were harvested in paraffin. IHC was performed as described (Hamon et al., 2019). Primary and secondary antibodies are listed in Supplementary Table 1. Sections were counterstained with 1:1000 4′,6-diamidino-2-phenylindole (DAPI) (1 mg/mL, Thermo Scientific).

### EdU labeling and TUNEL assay

For EdU labelling, females were injected intraperitoneally with 10 mM of 5-ethynyl-20-deoxyuridine (EdU) (Life Technology). EdU incorporation was detected on paraffin sections or frozen sections using the Click-iT EdU Imaging Kit following manufacturer’s recommendations (Life Technology). Apoptosis was detected by terminal deoxynucleotidyl transferase-mediated biotinylated UTP nick end labeling (TUNEL) assays using *in situ* cell death detection kit (Promega). All images were acquired using a Zeiss LSM710 confocal microscope and Zen software (Zeiss).

### Immunoblotting

Frozen retinas were lysed by sonication in lysis buffer (20 mM Na_2_HPO4, 250 mM NaCl, 30 mM NaPPi, 0.1% NP40, 5mM EDTA, 5mM DTT) supplemented with protease inhibitor cocktail (Sigma-Aldrich). Lysates concentration was determined using a Lowry protein assay kit (Bio-Rad) following sonication and centrifugation. Protein supernatants were separated under denaturing condition by SDS-PAGE, transferred onto nitrocellulose membrane and probed with antibodies (listed in Supplementary Table 1.), as described (Hamon et al., 2019). Proteins were visualized using enhanced chemiluminescence kit (Bio-Rad). α-tubulin was used for normalization. Quantification was performed using ImageJ software (http://imagej.nih.gov/ij/; provided in the public domain at NIH).

### Retinal flat mount

Fixed retinas were permeabilized and blocked in a solution containing 0.5% Triton-X100, 5% donkey normal serum, 1XPBS, 0.1 g/L thimerosal for 1 day at RT under agitation. Primary antibodies were diluted in a solution containing 0.5% Triton-X100, 5% donkey normal serum, 10% Dimethyl Sulfoxide, 1XPBS, 0.1 g/L thimerosal for 3 days at RT under agitation. The retinas were then washed for 1 day in PBST (1XPBS, 0.5% Triton-X100). Secondary antibodies were diluted in the same solution as primary antibodies and left for 2 days. After washing retinas for 1 day, they were mounted on slides and imaged using a scanning confocal microscope (Olympus, FV1000). Primary and secondary antibodies are listed in Supplementary Table 1.

### Electroretinography

Electroretinogram (ERG) recordings were performed using a focal ERG module attached to Micron IV (Phoenix Research Laboratory). Briefly, mice were dark-adapted overnight and prepared for the experiment under dim-red light. The mice were anesthetized with ketamine (100 mg/kg) and xylazine (10mg/kg) and received topical proparacaine hydrochloride (0.5%, Alcon) via eye drops. Pupils were dilated with tropicamide (1%, Alcon) and phenylephrine (2.5%, Alcon) and lightly coated with GONAK hypromellose ophthalmic demulcent solution (2.5%, Akorn). Lens of the Micron IV was placed directly on the cornea, and a reference electrode was placed on the mouse head. Scotopic responses were elicited with a series of flashes of increasing light intensities from −1.7 to 2.2 cd.s/m^2^. Photopic responses were elicited under rod-desensitizing background light with a series of flashes of increasing light intensities from −0.5 to 2.8 cd.s/m^2^. Values of a- and b-wave were extracted and plotted for comparisons between groups of interest.

### Optomotor response (OMR)

Real time video tracking and automated measurements of compensatory head movements in freely moving mice were performed using an OMR recording setup (Kretschmer et al., 2013, 2015) (Phenosys, Berlin, Germany). Each mouse was placed on a platform in the center of four computer-controlled LCD monitors. Visual stimuli were sinusoidally modulated luminance gratings generated by four LCD screens (60 Hz refresh rate; OkrArena, PhenoSys GmbH, Berlin, Germany), presented with a constant rotation. Video tracking considered the animal’s distance to the monitors, thereby keeping the spatial frequency of the retinal image constant and providing data for automated OMR quantifications.

OMRs were recorded using two different Michelson contrasts and different spatial frequencies (presented in random order) in the two mouse groups: 100% contrast or 50% contrast (n=13 for *Gsk3α^f/+^β^f/f^* and 18 for *Gsk3α^f/+^β^f/f^*;*α-Cre* genotype). All stimuli were presented for 60 s randomly in either clockwise or counterclockwise direction. The measurements were completed in three trials for each animal. At 100% and 50% contrast, OMRs were recorded in response to sinusoidal gratings at 12 spatial frequencies between 0.0125 and 0.5 cycles per degree (cpd). The number of head movements recorded at a speed range from 2 to 14 degrees per second in the same direction as the stimulus (T_Correct) and in the opposite direction (T_Incorrect) were used to calculate the OMR indices (T_Correct / T_Incorrect) at each spatial frequency.

### Retrograde labeling of retinal ganglion cells

For retrograde labeling, eyes were enucleated with a piece of the optic nerve and fixed in 4% PFA for 30 min. Rhodamine B isothiocyanate–Dextran (Sigma-Aldrich) was applied on the top of the optic nerve and incubated for 60 min. Eyes were flat mounted after the remaining dye was washed out for 48 hours in PBS at 4°C. Z series images were acquired using SP5 confocal microscope (Leica Biosystems), and 3D reconstruction was performed using Volocity (Perkin Elmer).

### Anterograde labeling of retinal ganglion cell projections

#### Anterograde labeling

For anterograde tracing of retinal projections, a Cholera Toxin beta subunit (CTB) was used. Animal were anesthetized using a cocktail of ketamine (60 mg/kg) and xylazine (10 mg/kg) and a subsequent bilateral injection of 1.2uL CTB at 1mg/ml coupled to either an Alexa-555 or -647 (Lifesciences) were performed intravitreally. Three days following the injection, mice were perfused with 4% PFA.

#### Tissue Clearing and 3D imaging

For 3D imaging of CTB-labeled brains, a methanol clearing protocol was carried out using modification from the iDISCO+ protocol (Belle et al., 2014, 2017). Briefly, brains were dehydrated by immersion in progressive baths of methanol/1X PBS (20%, 40%, 60%, 80%, 100%, 100%) for 2 hours each at RT on a tube rotator (SB3, Stuart) at 14rpm, using a 15 mL centrifuge tube (TPP, Dutcher) protected from light. Following these baths, samples were immersed overnight in 2/3 Dichloromethane (DCM; Sigma-Aldrich) and then a 30-min bath in 100 % DCM before being transferred in Di-Benzyl Ether (DBE; Sigma-Aldrich) overnight prior imaging.

3D imaging/Image acquisition for all samples was performed as previously described (Dobin et al., 2013). Acquisitions were done using an ultramicroscope I (LaVision Biotec) with the ImspectorPro software (LaVision Biotec). The step size between each image was fixed at 2 μm with a numerical aperture of 0.120 and 150ms acquisition using a PCO Edge SCMOS CCD camera (2,560 x 2,160 pixel size, LaVision BioTec).

#### Image analysis

Imaris x64 software (Version9.1.2, Bitplane) was used for all image analysis. Stack images were first converted from .tiff to .ims files using the Imaris file converter v9.1.2. 3D reconstruction was visualized with the “volume rendering” function. To isolate ipsi- and contralateral MTN volumes, manual segmentation was carried out using the “surface” tool and the isoline selection (density, 10%). Each ipsi- and contra-lateral projection of the MTN was segmented to generate a volume (μm^3^). Movie reconstruction with .tiff series were done with ImageJ (1.50e, Java 1.8.0_60, 64-bit) and iMovie (version 10.1.1).

### Whole transcriptome sequencing and data analysis

Whole transcriptome analysis was performed on three independent biological replicates from *Gsk3α^f/+^β^f/f^;α-Cre* and *Gsk3α^f/+^β^f/f^* retina at P60. After harvesting, both retinas for each animal were immediately frozen. RNA was extracted using Nucleospin RNA Plus kit (Macherey-Nagel). RNA quality and quantity were evaluated using a BioAnalyzer 2100 with RNA 6000 Nano Kit (Agilent Technologies). Stranded RNA-Seq libraries were constructed from 100 ng high-quality total RNA (RIN > 8) using the TruSeq Stranded mRNA Library Preparation Kit (Illumina). Paired-end sequencing of 40 bases length was performed on a NextSeq 500 system (Illumina). Pass-filtered reads were mapped using STAR and aligned to mouse reference genome GRCm38.94 (Dobin et al., 2013). Count table of the gene features was obtained using FeatureCounts (Liao et al., 2014). Normalization, differential expression analysis and FPKM (fragments per kilobase of exon per million fragments mapped) values were computed using EdgeR (Chen et al., 2015). An FPKM filtering cutoff of 1 in at least one of the 6 samples was applied. A False Discovery Rate (FDR) of less than or equal to 0.05 was considered significant and a fold change cutoff of 1.5 was applied to identify differentially expressed genes. Comprehensive gene list analysis, enriched biological pathways, gene annotation, were based on Gene Ontology classification system using Metascape (Zhou et al., 2019). Data visualization was done using GOplot R package (Walter et al., 2015). To evaluate the expression of the DEGs in RGCs, we used published whole transcriptome analysis from purified RGCs available on Gene Expression Omnibus database (GSE87647) (Sajgo et al., 2017).

### Gene expression analysis by Real-Time PCR (RT-qPCR)

After RNA extraction using Nucleospin RNA Plus kit (Macherey-Nagel), 500 ng of total RNA was reverse transcribed using the iScript cDNA Synthesis Kit according to manufacturer instructions (BioRad). Primers used for RT-qPCR are shown in Supplementary Table 2. For each RT-qPCR, 2 μL of a ten-fold dilution of synthetized cDNA was used, and the reactions were performed in technical triplicates on a C1000 thermal cycler (CFX96 real-time system, BioRad) using SsoFast EvaGreen Supermix (BioRad) as previously described (Livak and Schmittgen, 2001). RT-qPCR experiments were performed on three independent biological replicates. Differential expression was determined using the ΔΔCt method with the geometric average of *Rps26* and *Srp72* as endogenous controls (Livak and Schmittgen, 2001).

### Statistical analysis

Statistical analysis was performed with GraphPad Prism 8.3.0 (GraphPad Software, La Jolla California USA). Results are reported as mean ± SEM. Nonparametric Mann-Whitney U test was used to analyze cell counting and qPCR data. *P* value ≤ 0.05 was considered significant. For OMR assay statistical analysis, a Grubbs test was performed at 5% to remove outliers followed by 2-way ANOVA (genotype and spatial frequency) with Bonferroni Post Hoc test. *P* value ≤ 0.05 was considered significant.

## ACKNOWLEDGEMENT

We are grateful to Dr. Jim Woodgett for providing the *Gsk3* floxed lines and Tudor Badea and Peter Gruss for the alpha-Cre line. We would like also to thank Leah Thomas, Elodie-Kim Grellier, Yide Mi, and Jessica Gumerson for their help with mouse colonies and technical support. This research was supported by the CNRS, Retina France and by the Intramural Research Program of the National Eye Institute (EY000450 and EY000546). EK was supported by the Ernst Ludwig Ehrlich Studienwerk.

## AUTHOR’S CONTRIBUTION

E.K., R.J.V., C.H., designed and performed the experiments and analyzed the data, S.L. AND P.S. performed the experiments and analyzed the data, A.C. designed the experiments, analyzed the data and revised the manuscript, M.P., A.S., J.E.R. designed the study, analyzed the data, wrote the manuscript with the help of E.K. R.J.V and C.H. J.E.R supervised the study.

## SUPPLEMENTARY FIGURE LEGENDS

**Supplementary data 1.**
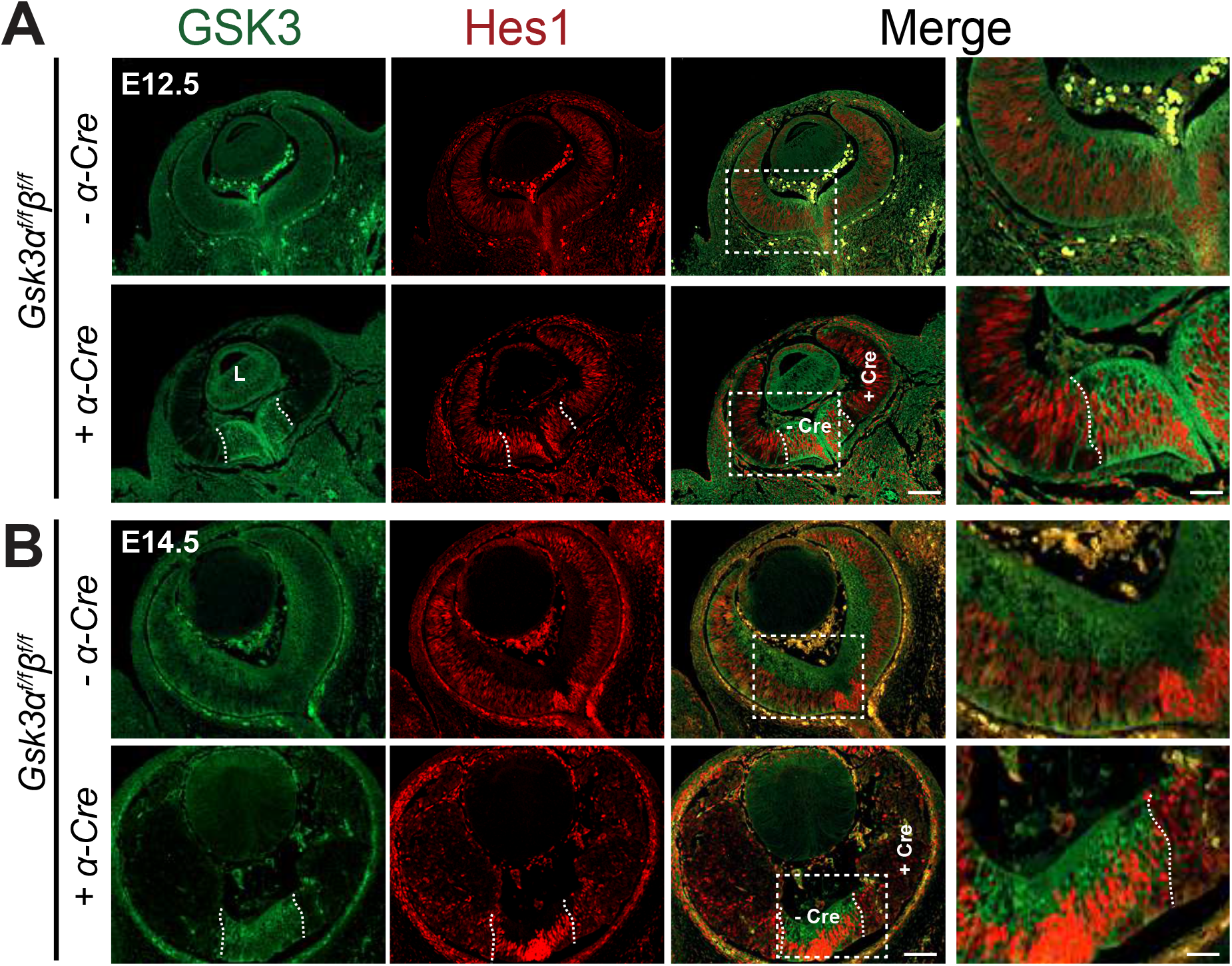
Lack of *Gsk3α* and *Gsk3β* expression impairs the maintenance of the pool of retinal progenitors. IHC on E12.5 and E14.5 *Gsk3α^f/f^β^f/f^* and *Gsk3α^f/f^β^f/f^;α-Cre* retina using Hes1 (red) and pan-GSK3 (green) indicates a loss of retinal progenitors at E14.5. Magnification on the right panel shows the squared delimited area. Dashed lines delimit the central *Gsk3*-expressing area and the *Gsk3*-deleted area at the periphery Scale bar: 100 μm, magnification area scale bar 40 μm.

**Supplementary data 2.**
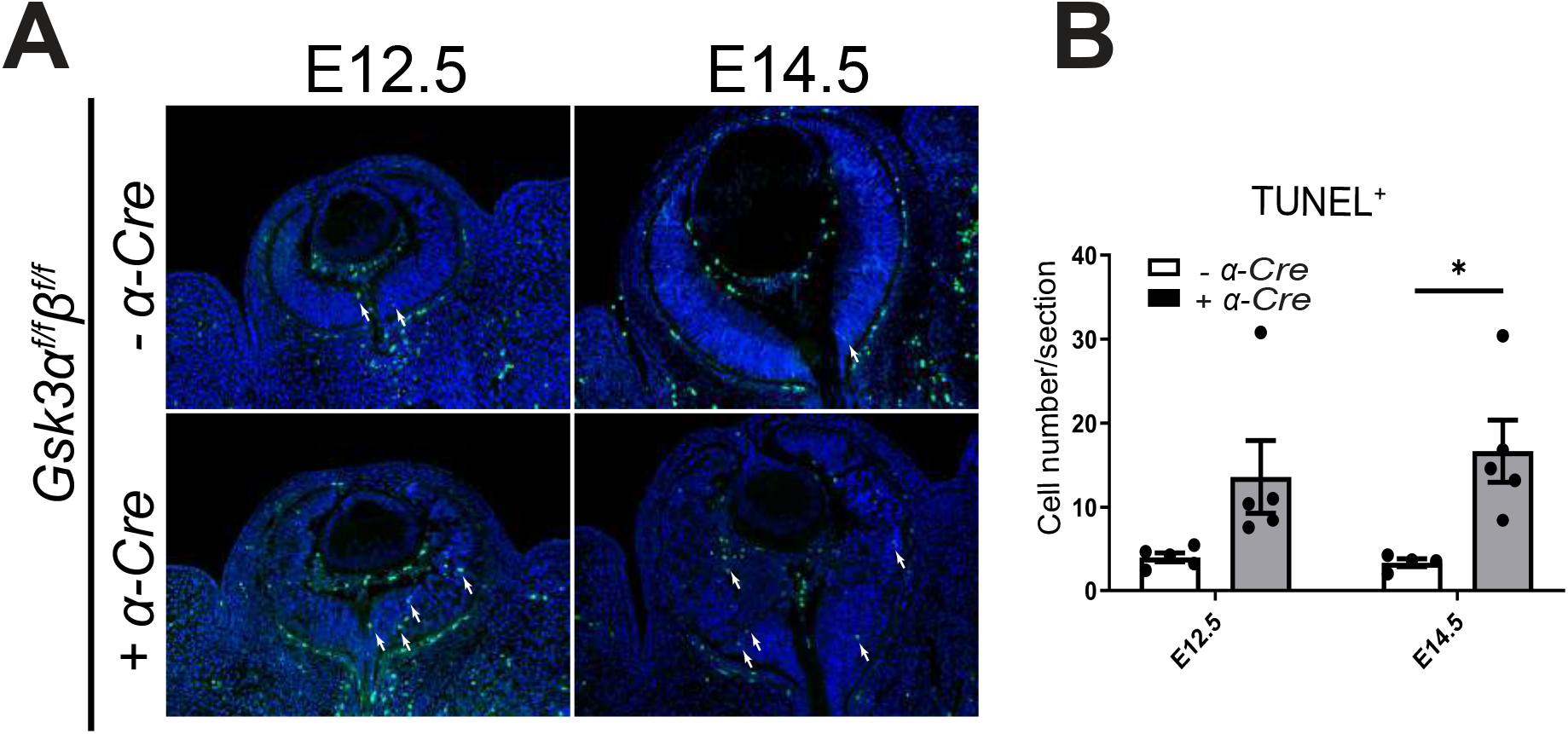
Lack of *Gsk3α* and *Gsk3β* expression in retinal progenitors leads to increased cell death. (A) TUNEL assay on E12.5 and E14.5 *Gsk3α^f/f^β^f/f^;α-Cre* and control animals reveals an increase of cell death (TUNEL-positive cells, green) in *Gsk3α^f/f^β^f/f^;α-Cre* animals compared to littermate controls. (B) Quantification of the number of TUNEL-positive cells per retinal section. Mean ± SEM values are presented from 5 to 6 biological replicates for E12.5, 4-5 biological replicates for E14.5, * indicates *P* ≤ 0.05.

**Supplementary data 3.**
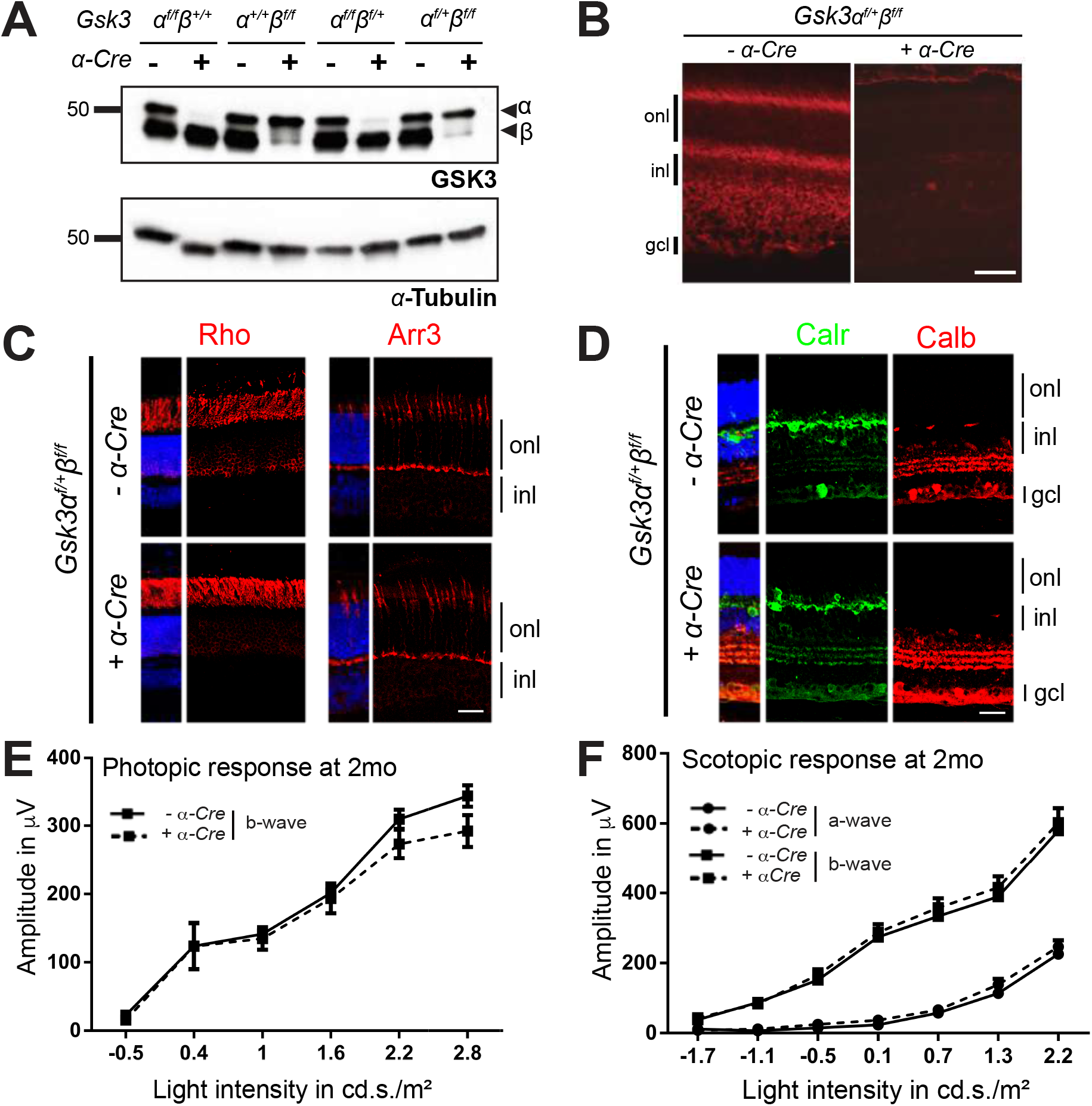
One allele of either *Gsk3α* or *Gsk3β* is sufficient for the development of a functional retina. (A) Immunoblot analysis of protein extracts from 2-month-old animals with different combination of *Gsk3α* and *Gsk3β* floxed alleles (*Gsk3α^f/f^β^+/+^, Gsk3α^+/+^β^f/f^*, *Gsk3α^f/+^β^f/f^* or *Gsk3α^f/f^β^f/+^*) with or without *Cre* recombinase using anti-panGSK3 antibody (recognizing both isoforms) reveals decreased expression of GSK3α or GSK3β (arrowheads). *α*-Tubulin is used as loading control. (B) IHC on 2-month-old retinal sections from control and *Gsk3α^f/+^β^f/f^*; *α-Cre* retinas with or without Cre recombinase using anti-GSK3β antibody (red) showing ubiquitous *Gsk3β* expression in all retinal layers, whereas its expression is lost in the Cre-expressing retina. (C) Expression of only one *Gsk3* allele (*Gsk3α* is sufficient for proper photoreceptor development. IHC using anti-Rhodopsin (Rho, red) and anti-Cone arrestin (Arr3, red) antibodies to label rod and cone photoreceptors, respectively. (D) Expression of only one *Gsk3* allele (*Gsk3α* is sufficient for proper interneuron development. IHC using anti-Calretinin (Calr, green) and anti-Calbindin (Calb, red) antibodies to label horizontal and amacrine cells, respectively. (B-D) onl, outer nuclear layer; inl, inner nuclear layer; gcl, ganglion cell layer. Scale bar: 20μm. (E, F) Electroretinogram (ERG) recording in 2-month-old *Gsk3α^f/+^β^f/f^;α-Cre* animals and littermate controls. Photopic (cones) (E) and scotopic (rods) (F) response in *Gsk3α^f/+^β^f/f^;α-Cre* animals are similar to controls. Mean ± SEM intensity response curves of a- and b-wave responses averaged from 8 biological replicates of each genotype.

**Supplementary data 4.**
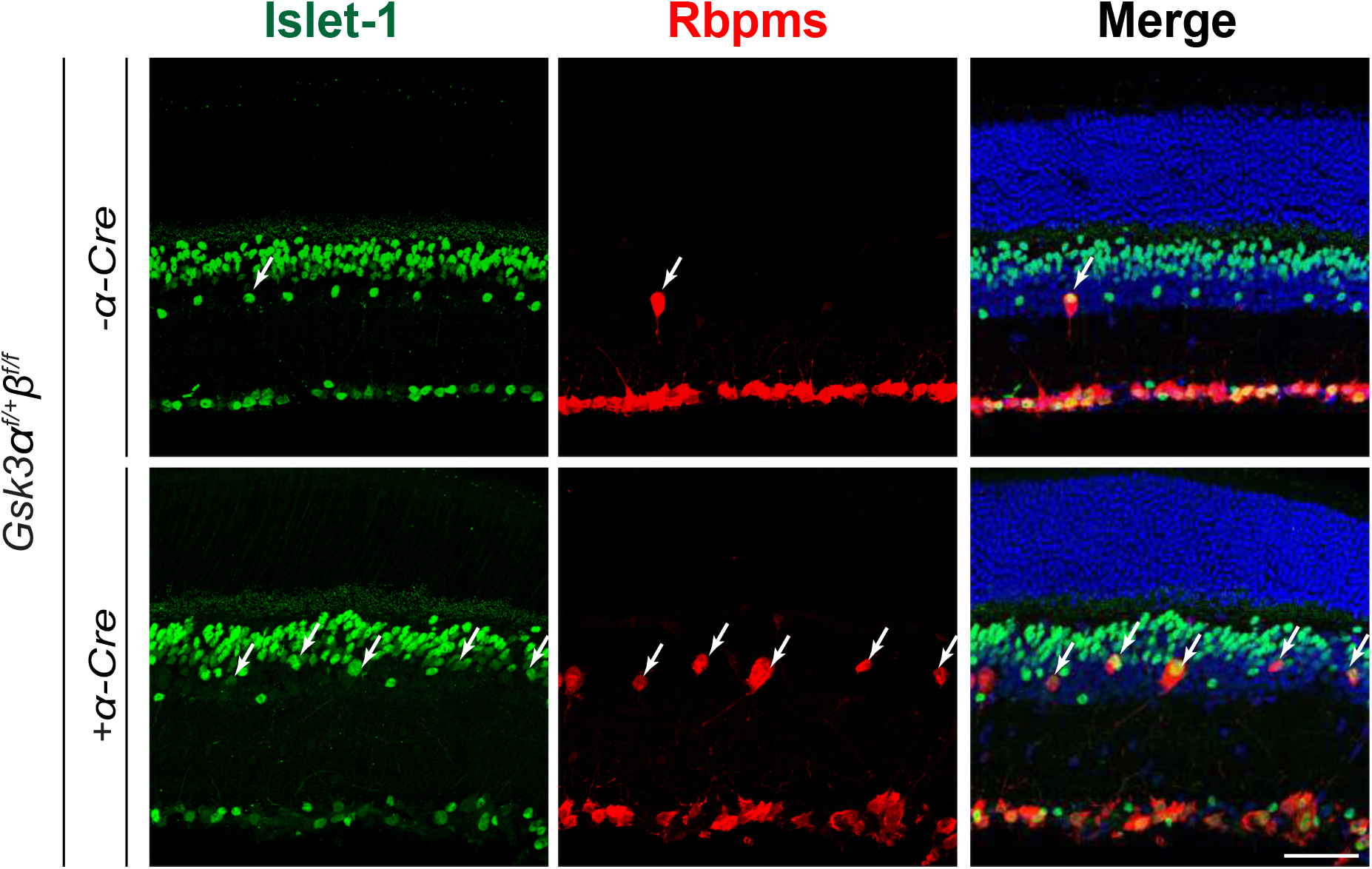
dRGCs express the nuclear factor Islet-1. IHC on 2-month-old mouse retina reveals that most dRGCs (Rbpms-positive dRGCs, white arrows, red) in the INL of *Gsk3α^f/+^β^f/f^; α-Cre* and littermate controls were positive for Islet-1 (green), a marker expressed in the nuclei of ganglion cells, and of cholinergic amacrine cells, ON-bipolar cells, and subpopulations of horizontal cells. Scale bar: 50 μm.

**Supplementary data 5.**
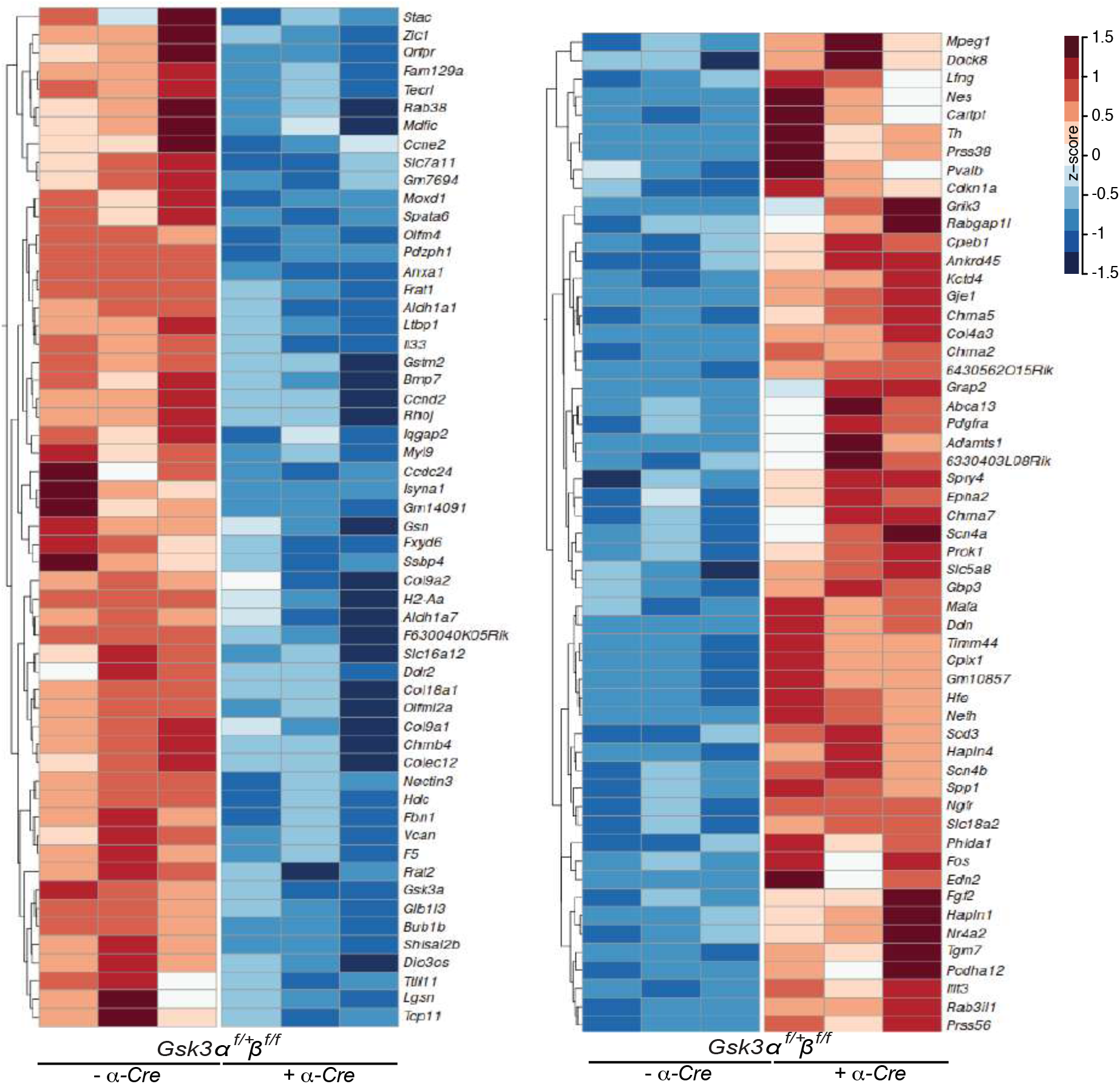
Hierarchical clustering of the identified differentially expressed genes. Hierarchical clustering representing the 111 DEGs (abs(FC)≥1.5; FDR≤0.05; FPKM>1) between 2-month-old *Gsk3α^f/+^β^f/f^; α-Cre* retina and littermate control were clustered by their Z-score. Left panel corresponds to downregulated genes; Right panel corresponds to upregulated genes.

**Supplementary data 6.**
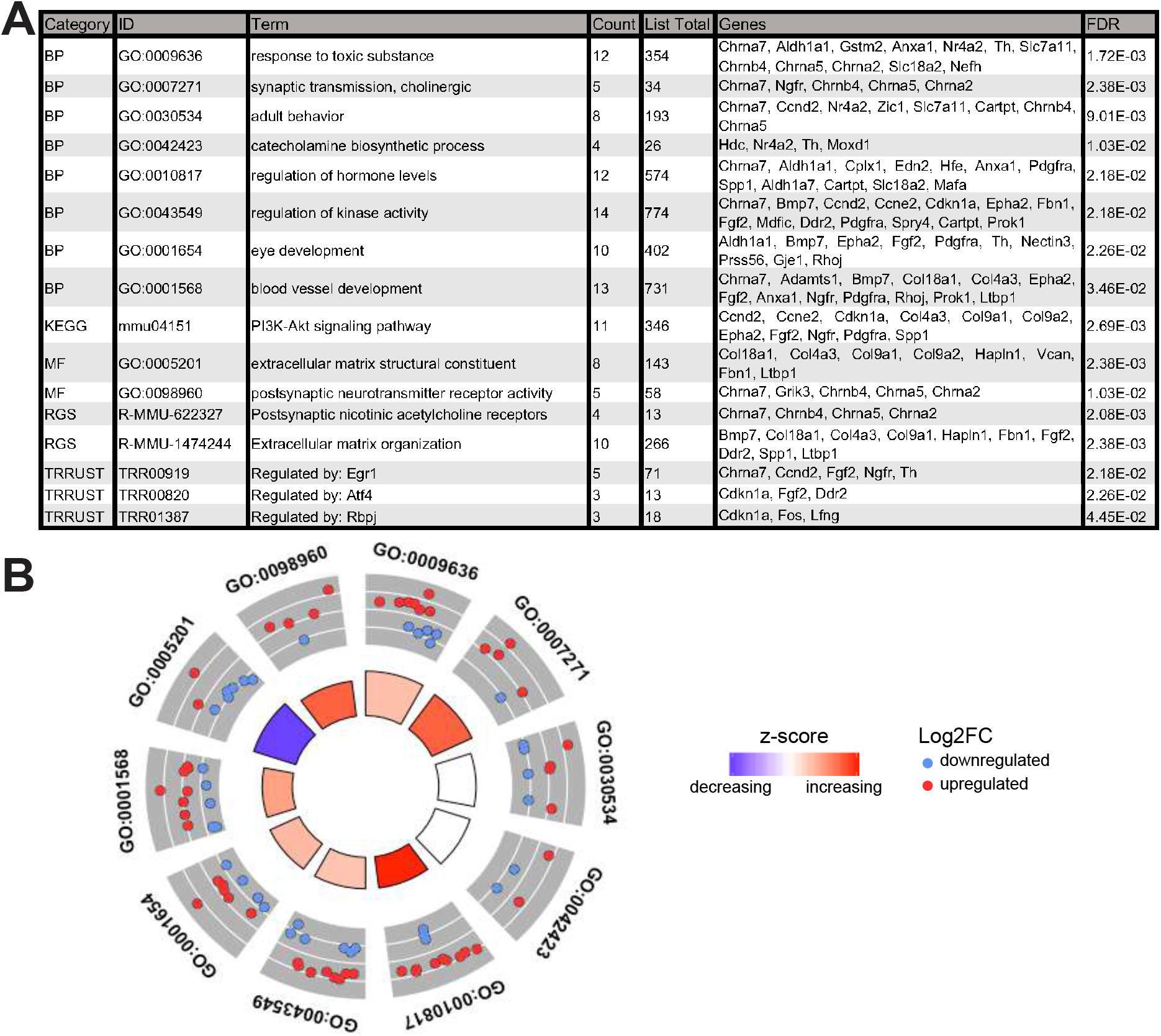
Identification of enriched pathways from DEGs identified in 2-month-old *Gsk3α^f/+^β^f/f^; α-Cre* retina. (A) Gene ontology (GO) annotations of DEGs in *Gsk3α^f/+^β^f/f^; α-Cre* retina compared to littermate controls. Top over-represented pathways for Biological process (BP), Molecular Function (MF), KEGG (Kyoto Encyclopedia of Genes and Genomes) and TRRUST (Transcriptional Regulatory Relationships Unrevealed by Sentence-based Text mining) were identified by enrichment analysis using Metascape. (B) Circular visualization for BP and MF of GO enrichment analysis. Down-regulated genes (blue dots) and up-regulated genes (red dots) within each GO pathway are plotted based on logFC. Z-score bars indicate if an entire GO category is more likely to be increased or decreased based on the genes within it.

**Supplementary Table 1.** List of primary and secondary antibodies used for immunohistochemistry (IF) and western blot (WB)

**Primary antibodies**
| ANTIGENE | HOST | SUPPLIER | REFERENCE | DILUTION (IF) | DILUTION (WB) |
| --- | --- | --- | --- | --- | --- |
| α-tubulin | mouse | SIGMA | T5168 | | 1:200.000 |
| GSK3α/$\beta$ | mouse | Thermo Fisher Scientific | 44-610 | 1:250 | 1:1000 |
| GSK3β | mouse | BD | 610201 | 1:250 | |
| β-catenin | rabbit | Abcam | ab2365 | 1:200 | |
| HES1 | rabbit | Cell Signaling | 11988S | 1:200 |  |
| Brn3a | mouse | Santa Cruz | sc-8429 | 1:200 |  |
| GFP | goat | Abcam | ab290-50 | 1:500 |  |
| pH3 | rabbit | EMD Millipore | 06-570 | 1:300 |  |
| Calbindin D-28k | rabbit | Swant | 300 | 1:100 |  |
| Calretinin | mouse | EMD Millipore | MAB1568 | 1:1000 |  |
| Cone Arrestin | rabbit | EMD Millipore | AB15282 | 1:1000 |  |
| Rhodopsin | Mouse | Abcam | MAB5316 | 1:2000 |  |
| Tbr2 | Rat | Ebioscience | 14-4876 | 1 :300 |  |
| Foxp2 | Goat | Santa Cruz | sc-21069 | 1 :1000 |  |
| Rbpms | Rabbit | PhosphoSolutions | 1830-RBPMS | 1 :400 |  |

**Secondary antibodies**
| ANTIGENE | HOST | SUPPLIER | REFERENCE | DILUTION (IF) | DILUTION (WB) |
| --- | --- | --- | --- | --- | --- |
| Alexa Fluor 555 anti-mouse IgG2A | goat | Thermo Fisher Scientific | A21127 | 1:1000 |  |
| Alexa Fluor 555 anti-mouse IgG2B | goat | Thermo Fisher Scientific | A21147 | 1:1000 |  |
| Alexa Fluor 488 anti-rabbit | donkey | Thermo Fisher Scientific | A21206 | 1:1000 |  |
| Alexa Fluor 488 anti-mouse IgG1 | goat | Thermo Fisher Scientific | A21240 | 1:1000 |  |
| Alexa Fluor 488 anti-rabbit | goat | Thermo Fisher Scientific | A21244 | 1:1000 |  |
| HRP anti-mouse IgG | goat | Sigma-Aldrich | A4416 |  | 1:5000 |

**Supplementary Table 2.**
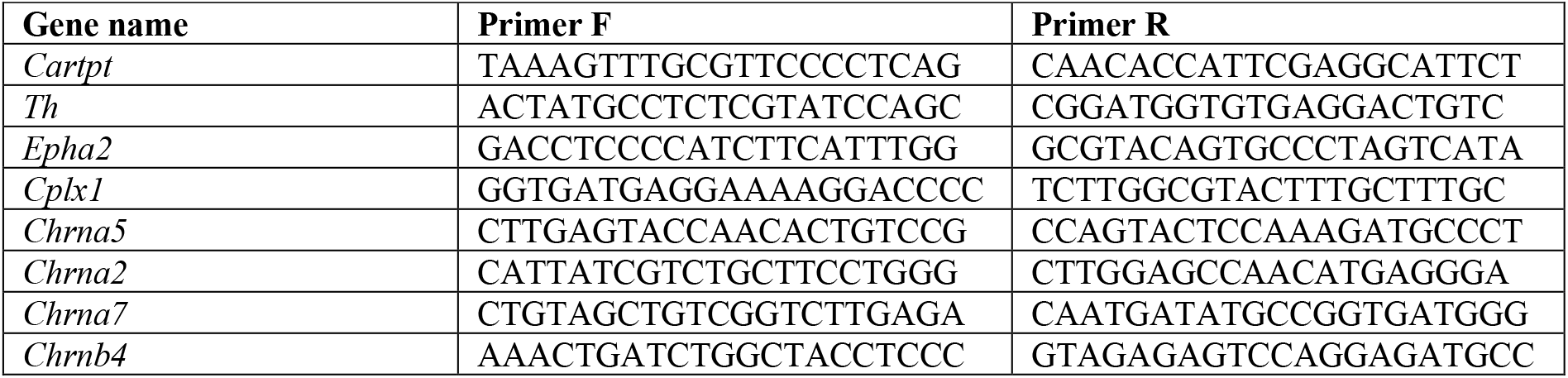
List of primers used for RT-qPCR.

